# Ovarian hormone deficiency enhances wood smoke-induced immune dysfunction via transcriptomic and metabolic alterations

**DOI:** 10.64898/2026.01.14.699504

**Authors:** Mijung Oh, Sydnee Yazzie, Eunju Lim, Onamma Edeh, Charlotte McVeigh, Alicia Bolt, Jennifer M. Gillette, Katherine E. Zychowski

## Abstract

The increasing frequency and severity of wildfires have heightened public exposure to smoke, highlighting the importance of identifying susceptibility factors, including ovarian hormone deficiency. Here, we used single-cell RNA sequencing to profile bone marrow immune cells from ovariectomized (OVX) mice exposed to either filtered air (FA) or wood smoke (WS) followed by functional validation in macrophages from both OVX and Sham-operated mice. Single-cell analyses focused on the OVX context; interactions between surgery and exposure were confirmed at the functional level in assays that included both Sham and OVX groups. In OVX mice, WS broadly suppressed transcriptional programs involved in antigen processing, leukocyte activation, antiviral defense, and bone remodeling. This was associated with altered immune cell composition, including increased memory CD8L T cells and decreased granulocytes and interferon-responsive populations. Bone marrow–derived macrophages (BMDMs) from WS-exposed OVX mice displayed metabolic reprogramming, characterized by the reversal of OVX-induced suppression of oxidative phosphorylation and glycolytic activity, along with reduced expression of M2-associated genes, without concurrent induction of M1-associated genes. This immune–metabolic decoupling suggests that WS exposure under ovarian hormone deficiency may imprint a lasting program in the bone marrow macrophage axis. Together, these findings show that ovarian hormone deficiency increases vulnerability to WS-induced immune disruption in the bone marrow. WS triggers macrophage reprogramming only under ovarian hormone deficiency, leading to heightened metabolic activity alongside suppression of key immune pathways, identifying a novel mechanism of immunotoxicity. These findings emphasize the need to consider hormonal status in air pollution risk assessment.

## Introduction

Wildfire smoke, a complex mixture of environmental toxicants, represents a significant public health challenge. Wildfires have become increasingly frequent and severe around the world, particularly in fire-prone regions such as the western United States (Agency 2024). This escalation is increasing human exposure to wood smoke (WS), a hazardous mixture of fine particulate matter (PM), volatile organic compounds (VOCs), and trace metals (Hadley et al. 2021). Inhalation of WS has well-documented acute and chronic health effects, especially on the respiratory system. Wildfire smoke exposure is linked to spikes in asthma and chronic obstructive pulmonary disease (COPD) exacerbations (Agency 2025; Reid et al. 2016; Wilgus and Merchant 2024) and to systemic immune perturbations characterized by widespread inflammatory signaling and dysregulation (Frumento and Țãlu 2024; Swiston et al. 2008). These systemic effects likely arise from circulating inflammatory mediators, indicating that the impact of inhaled toxicants extends beyond the lungs to distant organs and immune compartments.

While the pulmonary consequences of WS inhalation are widely recognized, its impact on the bone marrow, the central niche for immune cell development, remains poorly understood. The bone marrow is highly sensitive to environmental insults and plays a pivotal role in maintaining systemic immunity by continuously replenishing circulating immune cells (Zhao et al. 2012). A growing body of evidence from air pollution studies indicates that inhaled particulates can disrupt bone marrow functions; for example, fine particulate matter exposure can impair hematopoiesis, hinder progenitor cell mobilization, thus skewing the output of immune cells (Gangwar et al. 2020; Haberzettl et al. 2012). Such disruptions in the bone marrow compartment could lead to widespread immune dysregulation and increase susceptibility to inflammatory or immune-related diseases (Bhattarai et al. 2024; Mantovani et al. 2008). However, most wildfire smoke research has focused on the lungs and systemic circulation, with limited attention to the bone marrow’s role in mediating or propagating these immune effects. This knowledge gap is important because bone marrow–driven alterations may underlie long-term systemic consequences of air pollution exposure, including immune dysfunction and cancer susceptibility. This possibility aligns with emerging concepts of innate immune memory, or trained immunity, whereby environmental insults induce lasting epigenetic reprogramming of hematopoietic progenitors (Divangahi et al. 2021). Such maladaptive reprogramming could directly link acute exposure to the long-term establishment of a dysregulated immune state, contributing to chronic inflammation and impaired pathogen clearance (Saaoud et al. 2023).

Another critical but underexplored determinant of WS response is hormonal status. Sex hormones, particularly estrogen, modulate innate and adaptive immunity, contributing to sex differences in inflammation (Edwards et al. 2018; Rebuli et al. 2019). Estrogen generally dampens excessive pro-inflammatory signaling while supporting regulatory pathways (Lee and Chiang 2012; Markle and Fish 2014), so loss of ovarian hormones may dismantle a key checkpoint that constrains pollutant-induced inflammation. Consequently, hormone status is increasingly recognized as a modifier of pollutant-induced health outcomes (Commodore et al. 2024; Griffith et al. 2024). However, how ovarian hormone deficiency shapes immune responses to acute WS exposure remains largely unknown, despite its relevance for postmenopausal individuals and others with ovarian hormone loss, a large and biologically susceptible population (Gore et al. 2015). This has direct implications for refining public health risk assessment for airborne pollutants.

Experimental studies support the idea that ovarian hormone deficiency exacerbates pollutant-induced immune dysfunction. Animal models using ovariectomized (OVX) mice, a mouse model of ovarian hormone deficiency, have demonstrated heightened inflammation and immune dysregulation after pollutant exposure compared with hormonally intact controls (Adivi et al. 2022). Using the identical acute WS protocol as in our prior studies, we previously observed significant interactions between sex and treatment with evidence for female susceptibility at selected systemic and neuroinflammatory endpoints, and OVX females showed amplified inflammatory responses relative to Sham-operated mice (Wardhani et al. 2024a; Wardhani et al. 2024b). Despite these observations, the cellular and molecular mechanisms by which ovarian hormone deficiency conditions the response to WS remain unclear.

Here, we test the hypothesis that acute WS exposure in OVX mice would induce distinct transcriptomic and functional alterations in bone marrow immune progenitors and their macrophage progeny. To address this, we used a two-stage design: (i) single-cell discovery in OVX bone marrow to delineate the susceptibility-linked transcriptional landscape, followed by (ii) functional validation in bone-marrow–derived macrophages (BMDMs) from both Sham and OVX cohorts to formally test the interactions between surgery and treatment across metabolic and polarization-associated readouts. This framework allowed us to localize susceptibility within the myeloid compartment and to evaluate whether ovarian hormone deficiency is associated with macrophage functional dysregulation following acute WS exposure.

## Materials and Methods

### Animals and Surgical Procedures

Two separate cohorts of mice were used for these studies. For the single-cell RNA sequencing experiments, pre-ovariectomized (OVX) female C57BL/6 mice (n = 3 per group) were purchased at 6–8 weeks of age from The Jackson Laboratory (Bar Harbor, ME, USA; RRID: IMSR_JAX:000664). The cohort was intentionally restricted to mice to define the transcriptional landscape within the a priori most susceptible population, based on our prior work demonstrating significant ovarian hormone-dependent alterations in bone marrow immune cell populations following identical exposure (Wardhani et al. 2024b).

For the functional macrophage assays (metabolism and qRT-PCR), standard female C57BL/6 mice (n = 5 per group) were purchased at 6–8 weeks of age from Taconic Biosciences (Albany, NY, USA; RRID: IMSR_TAC: b6). This age range was selected due to its well-characterized immune profile, sexual maturity, and suitability for studying sex hormone effects on immune responses (Wardhani et al. 2024a; Wardhani et al. 2024b). Mice were housed under a 12-hour light/dark cycle with ad libitum access to food and water. To model postmenopausal estrogen deficiency, mice underwent bilateral ovariectomy. Anesthesia was induced using isoflurane (Sigma-Aldrich, Cat# 51144), and buprenorphine (Thermo Fisher Scientific, Cat# NC9122907) (0.01 mg/mL, 0.2 mL total) was administered subcutaneously for analgesia. The surgical site was sterilized with ethanol and betadine/iodine (Fisher Scientific, Cat# 19-066452), followed by a blunt dissection to expose and remove the ovaries. The fat pads were repositioned, and a single peritoneal stitch was used to close the incision. All mice were monitored for 10 days post-surgery to assess post-operative distress and incision healing.

To physiologically validate the effective contrast in hormonal status, body weight was recorded at two time points (pre-surgery baseline and immediately pre-exposure). We report weight gain as pre-exposure minus pre-surgery; OVX) mice showed greater weight gain than Sham (median 4.2 g vs 1.9 g; Mann–Whitney U=27, N_Sham_=19, N_OVX_=20, p<0.0001; Supplementary Figure S1). Sham females were maintained under natural cycling, however estrous stage was not determined on exposure days.

All procedures were approved by the University of New Mexico Institutional Animal Care and Use Committee (IACUC protocol #23-201389-HSC) and conducted in compliance with ARRIVE guidelines.

### Wood smoke exposure and characterization

Following a two-week acclimation, mice were assigned to a 2-day exposure regimen that included daily 4-h intervals of acute wood smoke (WS) at a concentration of approximately 0.575 ± 0.12 mg/m³, a setup that emulated an acute exposure to WS from a wildfire event. This concentration is consistent with real-world exposure levels, which were previously measured to be 0.5 mg/m³ (Nalesnik et al. 2025). WS was generated from 2 g piñon biomass and delivered via the same combustion/dilution/pressurization system described previously. Chamber gases (NO, CO, OL) were logged every 10 min using a Grey Wolf TG501, with detailed time-series results reported previously (Wardhani et al. 2024b), while chamber temperature was maintained at 21.3–21.5 °C.

The physical and chemical properties of WS generated by this identical system were previously characterized (Wardhani et al. 2024b) and are summarized here for reference. This prior characterization determined the WS particle size by Aerosizer (MMD 0.13 μm and GSD 1.4; Supplementary Table S1) and assessed particle-phase metals by ICP-MS, which showed significant elevations in L³Cu, ¹L²W, ²LLPb, ²³LU compared with FA filters (Supplementary Table S2). All instrumentation and QA/QC matched this prior characterization. The control condition involved exposure to HEPA-filtered air (FA) in a chamber for the same timeframe. The same whole-body WS protocol used here has been previously shown to induce a clear systemic plasma lipidomic signature in female mice, with significant decreases in multiple phosphatidylcholines and phosphatidylethanolamines by targeted LC-MS lipidomics, supporting the biological effectiveness of this exposure paradigm at the organismal level. We cite those data here as prior validation for the present study; the plasma analysis was sex-stratified (female vs. male) but not further stratified by OVX status (Wardhani et al. 2024a).

### Bone marrow-derived immune cell collection

Mice were euthanized 24 hours after the final WS or FA exposure. Both femurs and tibias were excised under sterile conditions and cleaned of muscle tissue using gauze soaked in 70% ethanol. The bones were placed in cold RPMI-1640 (Gibco, Cat# 11875085) medium supplemented with 2% fetal bovine serum (Gibco, Cat# 10438026) and kept on ice until processing.

Bone marrow was extracted by inserting a 25-gauge needle attached to a 10 mL syringe filled with cold RPMI + 2% FBS and flushing the marrow cavity of each bone into a sterile 15 mL conical tube. Flushed marrow from each mouse was gently triturated with a pipette to dissociate cell aggregates and obtain a single-cell suspension. The resulting cell suspensions were passed through a 70 μm cell strainer (Corning, Cat# 352350) into clean tubes to remove bone fragments and debris.

Cell viability and concentration were assessed using the Nexcelom Cellometer Auto2000 (Nexcelom Bioscience) with acridine orange/propidium iodide (AOPI) fluorescent staining. Viable bone marrow–derived immune cells were collected for downstream applications. To assess the transcriptional impact of WS exposure under ovarian hormone–deficient conditions, scRNA-seq was performed on samples from OVX mice exposed to either FA or WS.

### Library Preparation and Single-Cell RNA Sequencing

Sixteen thousand cells were loaded into the Chromium iX Controller (10X Genomics, PN-1000328) on a Chromium Next GEM Chip G (10X Genomics, PN-1000120) and processed to generate single-cell Gel Beads-in-Emulsion (GEMs) according to the manufacturer’s protocol. Each encapsulated cell was lysed within its GEM, releasing messenger RNA, which was reverse transcribed into cDNA using primers attached to the gel beads. Each gel bead carried a unique 10x barcode for downstream cell separation.

After reverse transcription, the GEMs were broken, and all uniquely barcoded cDNAs were pooled and PCR amplified to generate sufficient material for Illumina sequencing. cDNA and library construction were performed using the Chromium Next GEM Single Cell 3’ Reagent Kits v3.1 (10X Genomics, PN-1000286) and Dual Index Kit TT Set A (10X Genomics, PN-1000215) following the manufacturer’s manual. Quality control for the constructed library was conducted using the Agilent Bioanalyzer High Sensitivity DNA kit (Agilent Technologies, Cat# 5067-4626) for qualitative analysis and the Qubit DNA HS assay kit (Thermo Fisher Scientific, Cat# Q32851) for quantitative analysis.

The multiplexed libraries were pooled and sequenced on the Illumina NovaSeqX Plus platform at a depth of approximately 30,000 reads per cell. Sequencing was performed using 100-cycle kits with the following read lengths: 28 bp for Read1 (cell barcode and UMI) and 90 bp for Read2 (transcript).

### Bioinformatic Analysis of scRNA-seq Data

Raw sequencing data, in base call format (.bcl) was demultiplexed using Cell Ranger from 10x Genomics, converting the raw data into FASTQ format. Cell Ranger was also used for alignment of the FASTQ files to the mouse reference genome (mm39) and count the number of reads from each cell that align to each gene. The resulting matrix files which summarize the alignment results were imported in the Seurat package (v4.3 or later) (Satija Lab, NYGC) for further analysis.

In Seurat, each individual sample was preprocessed, normalized, and scaled. As part of quality control, the distribution of counts (UMIs), features (genes), and mitochondrial gene percentages in single cells for each sample was evaluated. Violin plots showing the distribution of features (Supplementary Figure S2A) and percent mitochondrial genes (Supplementary Figure S2B) were used to assess sample quality. Cells with high expression of either counts or features were excluded from further analysis, as these can indicate doublets or multiplets. Additionally, cells with high percent mitochondrial genes were also removed. All samples were combined into a single dataset, adding metadata with the original sample information. This combined dataset was evaluated for differences between samples and/or clusters using Seurat, including finding biomarkers (to help identify cell types) and performing differential expression comparisons between specific clusters. Dimensionality reduction was performed using principal component analysis (PCA), followed by Uniform Manifold Approximation and Projection (UMAP). Graph-based clustering was used to identify immune cell populations, which were annotated based on canonical marker gene expression. For the analysis of cell type proportions, a generalized linear mixed-effects model (GLMM) with a binomial distribution (logistic regression) was used to rigorously test for compositional changes while accounting for inter-animal variability. In this model, ‘Treatment’ (FA vs. WS) was included as a fixed effect, while ‘Biological Replicate’ (mouse ID) was included as a random effect. Odds ratios and p-values were calculated to determine the likelihood and significance of compositional shifts.

To assess the impact of WS exposure on bone marrow immunity, differential gene expression (DGE) analysis was performed across all immune cell populations (for Figure 2) and at the individual immune cell cluster level (for Figures 4 and 5), comparing FA- and WS-exposed group. For all DGE analyses, including the global transcriptomic view (Figure 2) and cluster-specific comparisons (Figures 4 and 5), DGE was determined using Seurat’s FindMarkers function. Statistical significance was assessed using FDR-adjusted p-values (Benjamini–Hochberg false discovery rate correction) with an FDR threshold of < 0.05 and |logLFC| > 0.5 as cutoffs. This rigorous approach formed the basis of our comprehensive analytical strategy: (1) cell-level pathway analysis incorporating effect size (median logLFC with 95% CI) visualization (Figure 4 and Figure5), (2) validation at the animal-replicate level using pseudobulk pathway analysis (Supplementary Figure S5), and (3) unbiased, orthogonal validation using Gene Set Enrichment Analysis (GSEA) (Supplementary Figure S6). For effect size visualization of these key cell-level DEGs, median log2 fold change (log2FC) and 95% confidence intervals (CI) were calculated. These effect sizes and CIs were visualized as forest plots for enhanced interpretability (Figure 4C, 4E, 5C, 5E, Supplementary Figure S4).

For pseudobulk pathway analysis, used to validate findings at the animal-replicate level, pseudobulk differential expression analysis was performed. For each cluster (Memory CD8+ T cells, ISG-expressing cells), gene counts were aggregated (summed) across all cells for each biological replicate (n=3/group).

Differential expression analysis was performed using standard RNA-seq analysis packages. To overcome the inherent low statistical power of n=3 vs. n=3 comparisons, a relaxed statistical threshold (unadjusted *p*-value < 0.05 and |logLFC| > 0.5) was used to generate a comprehensive candidate gene list for downstream pathway analysis. This gene list was then analyzed using Metascape (metascape.org) against GO and Reactome databases. A curated subset of the most relevant pathways is visualized in Supplementary Figure S5, while the complete list of all significant pathways (including the top 20 lists by *p*-value) and the full gene lists used for this analysis are provided in Supplementary Table S6.

For Gene Set Enrichment Analysis (GSEA) as a second, orthogonal validation method, GSEA was performed. This analysis does not rely on pre-selected DEG cutoffs. For each cluster, all genes were ranked based on their cell-level differential expression metric. GSEA was performed against the GO Biological Process database (c5.go.bp.v2023.Mmu). Pathways with a FDR *q*-value < 0.01 were considered highly significant. A curated summary plot of key pathways is presented in Supplementary Figure S6, and the full lists of all significant pathways are provided in Supplementary Table S7.

Data visualization included volcano plots generated in GraphPad Prism (v10.4) and heatmaps created using Morpheus. Functional enrichment analyses for Gene Ontology (GO) and Kyoto Encyclopedia of Genes and Genomes (KEGG) pathways were performed using Metascape to identify biological processes associated with WS-induced transcriptional changes.

### Bone Marrow-Derived Macrophage (BMDM) Differentiation

Bone marrow cells were isolated from femurs and tibias of mice 24 hours after exposure to either FA or WS, as described above. Red blood cells were lysed using ACK lysing buffer (Quality Biological, Cat# 118-156-101), and the remaining nucleated cells were washed and resuspended in macrophage differentiation medium consisting of RPMI supplemented with 10% FBS, 1% penicillin-streptomycin (Gibco, Cat# 15140122), and 50 ng/mL recombinant mouse M-CSF (PeproTech, Cat# 315-02). Cells were plated at a density of 5 × 10L cells per well in 15 cm petri dish and incubated at 37°C with 5% COL. Culture medium was replaced on day 3 and day 6. On day 7, adherent BMDMs were detached for downstream analysis. To detach BMDMs without damaging surface markers or affecting metabolic activity, cells were gently washed with PBS and incubated with 2 mM EDTA (Thermo Fisher, Cat# 15575-038) in PBS (prepared by dilution of 0.5 M EDTA stock solution) for 10 minutes at 37°C. Cells were then lifted using gentle pipetting and collected by centrifugation at 300 × g for 5 minutes. The harvested BMDMs were resuspended in appropriate assay buffer for downstream analyses.

### Flow Cytometry Validation of Macrophage Phenotype

After 7 days of M-CSF–driven differentiation on non–tissue-culture–treated petri dishes (Day 7), BMDMs were harvested to assess baseline phenotype prior to re-plating. A paired set of cultures was then re-seeded onto tissue-culture–treated plates and allowed to adhere for 24 h (Day 8) to match the substrate/conditioning used for functional assays; cells were subsequently harvested for a second phenotyping time point. Single-cell suspensions were incubated with an Fc block (BioLegend, Cat# 101319) and then stained with fluorophore-conjugated antibodies against CD11b (PerCP-Cy5.5, clone M1/70; Cell Signaling, Cat# 85601; 1:80 dilution) and F4/80 (violetFluor 450, clone BM8.1; Cell Signaling, Cat# 40781; 1:40 dilution). Corresponding isotype controls (IgG2b PerCP-Cy5.5®, Cat# 79201; IgG2b violetFluor™ 450, Cat# 52572; Cell Signaling) were used to establish background fluorescence and guide gating. After staining, cells were washed and resuspended in PBS with 2% FBS. Flow cytometry was performed on a Beckman Coulter CytoFLEX S, and ≥10,000 live singlet per sample were acquired. Data were analyzed using FlowJo software. Live single cells were gated based on FSC and SSC, and the proportions of CD11bL, F4/80L, and CD11bLF4/80L double-positive cells were calculated. In line with maturation-dependent behavior of in-vitro M-CSF BMDMs, F4/80 increased after 24 h adherence on the tissue-culture plates used for functional assays (Supplementary Figure S8; CD11bLF4/80L 42.7% on Day 7 → 69.1% on Day 8; >99% CD11bL among live singlets).

### Metabolic Flux Analysis

Bone marrow–derived macrophages were seeded at 80,000 cells per well in Seahorse XF96 cell culture microplates and incubated overnight in macrophage differentiation medium. For mitochondrial stress testing, cells were washed and incubated in Seahorse XF Base Medium supplemented with 10 mM glucose, 1 mM pyruvate, and 2 mM glutamine, adjusted to pH 7.4. Oxygen consumption rate (OCR) was measured using the XF Cell Mito Stress Test Kit (Agilent, Cat# 103015-100), with sequential injections of oligomycin (1 μM), FCCP (1 μM), and rotenone/antimycin A (0.5 μM each). For glycolysis stress testing, cells were washed and incubated in Seahorse XF Base Medium supplemented with 2 mM glutamine only, following the manufacturer’s protocol. Extracellular acidification rate (ECAR) was measured using the XF Glycolysis Stress Test Kit (Agilent, Cat# 103020-100), with sequential injections of glucose (10 mM), oligomycin (1 μM), and 2-deoxy-D-glucose (2-DG, 50 mM). All measurements were conducted on a Seahorse XF96 Analyzer. To normalize for variations in cell number, total protein content was quantified from each well immediately following the assay using a standard BCA assay. A protein-derived normalization factor was then calculated for each well as its protein content relative to the mean protein content of the Sham-FA control group and entered into the normalization matrix, such that the resulting OCR and ECAR values are linearly proportional to OCR and ECAR per total protein content. Data were analyzed with Seahorse Wave software (Agilent).

### Quantitative Real-Time PCR (qRT-PCR)

Total RNA was extracted from BMDMs using the RNeasy Mini Kit (Qiagen, Cat# 74104), and 2000 ng of RNA was reverse-transcribed using the High-Capacity cDNA Reverse Transcription Kit (Thermo Fisher Scientific, Cat# 4374966) according to the manufacturer’s instructions. Using iTaq universal SYBR Green Super Mix (Bio-Rad, Cat# 1725121), qRT-PCR was performed on a CXF OPUS 96 Real-Time PCR System (Bio-Rad). Pre-designed and validated QuantiTect Primer Assays (Qiagen) were used to measure the expression of the following target genes: *Arg1* (Genbank: NM_007482.5, Cat: QT00134288), *Ccl2* (Genbank: NM_011333.3, Cat: QT00167832), *Ccl5* (Genbank: NM_013997.3, Cat: QT01747165), *Il10* (Genbank: NM_010548.2, Cat: QT00106169), *Mrc1* (Genbank: NM_008625.2, Cat: QT00103012), *Pparg* (Genbank: NM_011146.4, Cat: QT00100296), *Tgfb1* (Genbank: NM_011577.2, Cat: QT00145250), *Cxcl1* (Genbank: NM_008176.3, Cat: QT00115647), *Il1b* (Genbank: NM_008361.4, Cat: QT01048355), and *Tnfa* (Genbank: NM_013693.3, Cat: QT00116564). Gene expression was normalized to the reference gene *18S rRNA* (Genbank: NR_003278.3, Cat: QT02448075). The use of 18S rRNA as a single reference gene was empirically validated. The expression stability of four candidate genes (18S rRNA, GAPDH, Actb, and Hprt1) was compared across all samples using the RefFinder analysis tool. This analysis confirmed that 18S rRNA was the most stable reference gene for normalization in our experimental system (Supplementary Figure S9). Reactions were run in technical triplicates, and expression levels were normalized to 18S rRNA using the ΔΔCt method (Livak and Schmittgen 2001).

### Western Blot Analysis

To determine if these transcriptional changes translated to altered protein levels, we performed Western blot analysis on BMDM lysates. BMDM cell pellets collected after differentiation were lysed on ice using RIPA buffer supplemented with protease and phosphatase inhibitor cocktails (Thermo Fisher Scientific). Lysates were cleared by centrifugation, and protein concentration was determined by BCA assay (Thermo Fisher Scientific). Equal amounts of protein (30 µg) were mixed with Laemmli sample buffer, heated at 70°C for 10 min, and resolved on 4-20% Mini-PROTEAN TGX Precast Gels (#4561094, Bio-Rad). Proteins were transferred onto PVDF membranes (#IPVH00010, Bio-Rad). Membranes were blocked with 5% BSA in TBST for 1 hour at room temperature and incubated overnight at 4°C with primary antibodies against ARG1 (#93668, Cell Signaling Technology, 1:1000), PPARγ (#2443, Cell Signaling Technology, 1:1000), or Vinculin (#13901, Cell Signaling Technology, 1:1000) diluted in 5% BSA/TBST. After washing with TBST, membranes were incubated with HRP-conjugated secondary antibody (#7074S, Cell Signaling Technology, 1:2000) for 1 hour at room temperature. Bands were visualized using the SuperSignal^TM^ West Femto Maximum Sensitivity Substrate (#34095, Thermo Fisher Scientific) on a ChemiDoc Imaging System (Bio-Rad). Band intensities were quantified using Image Lab software (Bio-Rad), and target protein levels were normalized to Vinculin.

### Statistical Analysis

Unless otherwise specified, n denotes biological replicates (mice per group) and the experimental unit is the individual mouse. GraphPad Prism v10.4 was used for plotting and simple statistics unrelated to RNA-seq modeling. For comparisons between two groups, an unpaired two-tailed Student’s t-test was used. For multi-group comparisons, such as the metabolic (Figure 6) and protein expression results (Figure 8), a two-way ANOVA with Tukey’s multiple comparisons test were used to evaluate the main effects (e.g., surgery, treatment) and their interaction. For the analysis of qRT-PCR data (Figure 7 and Supplementary Figure S11), which exhibited significant inter-replicate variability inherent to *ex vivo* BMDMs differentiation, a linear mixed-effects model (LMM) was employed for each target gene. This statistical approach was chosen as LMMs are specifically designed to account for such batch effects by modeling the source of inter-replicate variation as a random effect (Espín-Pérez et al. 2018). In our model, ‘Surgery’ and ‘Treatment’ and their interaction were included as fixed effects, while ‘Biological Replicate’ was included as a random effect. All statistical tests were performed on the ΔCt values (prior to 2−ΔΔCt transformation for visualization). Post-hoc multiple comparisons were conducted to identify specific group differences following a significant interaction effect.

Consistent with previous single-cell sequencing studies, we used n=3 biological replicates per group, which has been demonstrated to be sufficient for detecting relevant phenotypic differences in similar experimental settings (Hammond et al. 2019; Marsh et al. 2022). To rigorously test the key hypotheses generated from this discovery-phase experiment, we utilized an expanded cohort of n=5 mice per group for all subsequent functional validation assays, including metabolic profiling and gene expression analysis. Animals were assigned to experimental groups based on surgical status (Sham or OVX) and randomized to FA or WS exposure. Investigators were not blinded during data collection, but data processing (e.g., clustering, qRT-PCR normalization) was performed using automated pipelines.

Data are presented as mean ± standard deviation (SD) unless otherwise indicated. Each dot in dot plots or bar graphs represents an individual animal unless otherwise noted. All statistical assumptions (e.g., normality of residuals) were tested using Prism’s built-in diagnostics prior to applying parametric methods. No data points were excluded from the analysis. Statistical results are explicitly shown in the figure panels, figure legends, or results text, and are summarized with *p*-value annotations as follows: ns = not significant, \**p* < 0.05, \*\**p* < 0.01, \*\*\**p* < 0.001, \*\*\*\**p* < 0.0001.

## Results

### Wood smoke exposure reshapes the bone marrow immune landscape in ovarian hormone-deficient mice

As illustrated in the experimental design (Figure 1A), OVX female mice were exposed to either WS or FA for two consecutive days. Exposure conditions matched our prior system performance (Wardhani et al. 2024b), after which bone marrow was collected for scRNA-seq profiling. After quality control and dimensionality reduction, Uniform Manifold Approximation and Projection (UMAP) clustering identified 16 distinct immune cell populations from the scRNA-seq data, including T cells, B cells, monocytes, neutrophils, dendritic cells, and progenitor cells (Figure 1B). Quality control metrics confirmed comparable sequencing depth and mitochondrial read percentages across all samples (SupplementaryLFigureLS2). Among the OVX female mice, comparative analysis between the FA and WS exposure groups revealed subtle but consistent shifts in bone marrow immune cell composition in OVX mice (Figure 1C), suggesting that WS alters the immune architecture under ovarian hormone-deficient conditions. These shifts, along with transcriptional suppression in granulocytes and interferon-stimulated gene (ISG)-expressing subsets, suggest that WS exposure may alter myeloid progenitor function, consistent with emerging concepts of innate immune memory (Kaufmann et al. 2018; Ochando et al. 2023). Cell type identity was defined based on the expression of canonical marker genes. For instance, major populations were identified as follows: Memory CD8L T cells (e.g., Fn1), granulocytes (Epx), neutrophils (Ltf), intermediate monocytes (Il1b), myeloid dendritic cells (Ace), and pro-B cells (Vpreb1), among others (Figure 1D).

**Figure 1.**
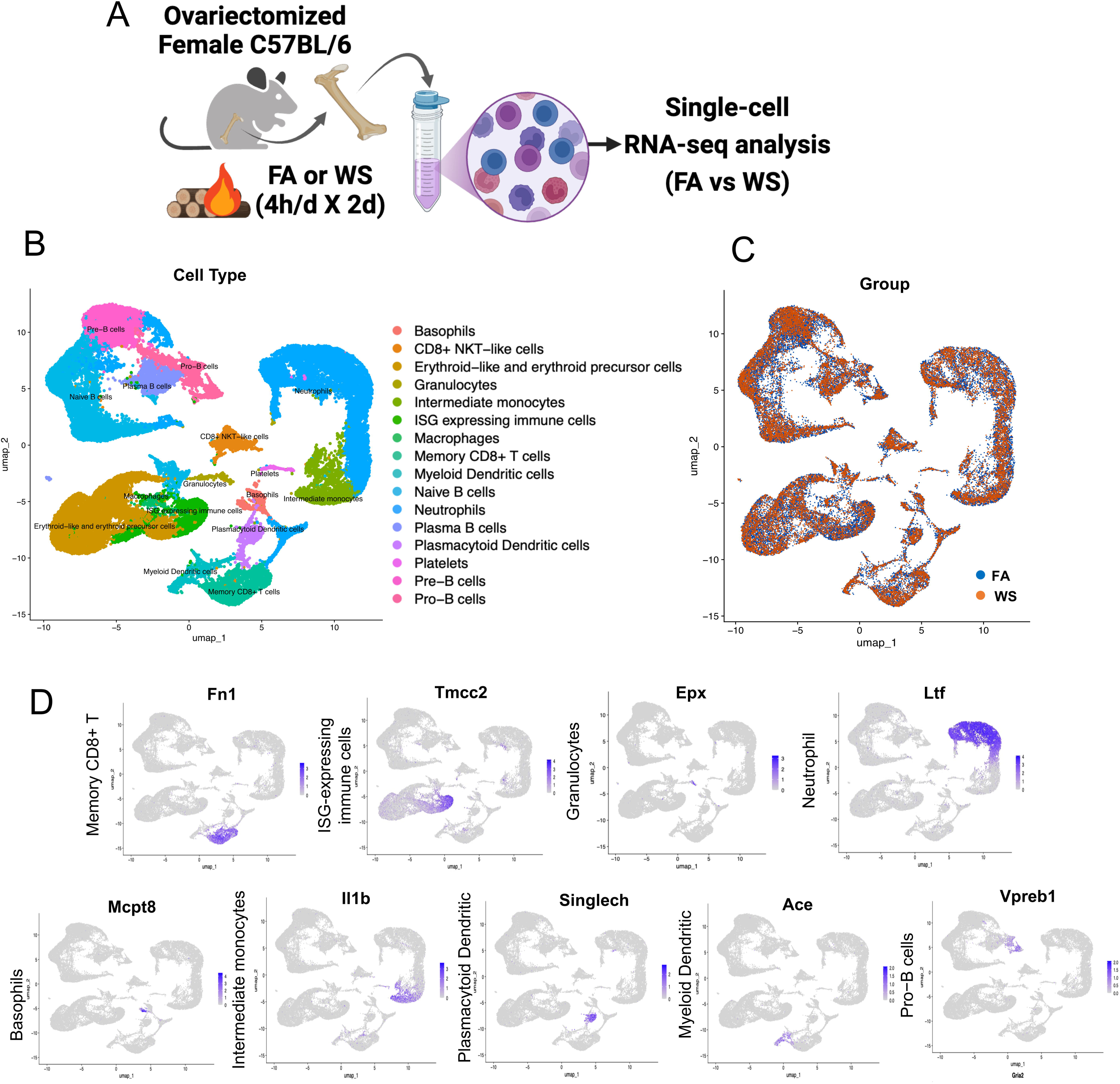
Experimental design and bone marrow immune cell composition in OVX mice after WS exposure. (A) Schematic of the experimental design, showing two exposure groups. Ovariectomized (OVX) female C57BL/6 mice underwent a 2-day exposure protocol consisting of daily 4-hour exposures to either acute wood smoke (WS) from wildfire combustion or control filtered air (FA) (n = 3 mice per group). All mice were euthanized 24 hours after the final exposure. (B) Uniform Manifold Approximation and Projection (UMAP) plot displaying the clustering of bone marrow immune cells from FA and WS exposure groups, revealing the major immune cell populations present. (C) UMAP plot showing the distribution of these immune cell clusters in FA vs. WS groups, indicating relative shifts in cluster abundance between the two conditions. (D) Feature plot illustrating the expression of key marker genes that define each of the major immune cell clusters identified in the dataset.

### Wood smoke broadly suppresses immune gene programs and alters immune composition in the bone marrow

To first understand the global impact of WS exposure, we analyzed the transcriptomic changes across all identified immune lineages. Across all immune lineages, 2,728 genes were significantly differentially expressed, with 528 showing an absolute logL fold change > 0.5 (Figure 2A). The vast majority of these (503 genes) were downregulated in WS-exposed OVX mice (Figure 2B). The suppressed genes spanned pathways involved in protein conjugation, antiviral defense responses, leukocyte differentiation, interleukin-12 production, Toll-like receptor signaling, and type I interferon response (Figure 2C), collectively indicating an impaired antiviral and pro-inflammatory capacity in the bone marrow under WS exposure. Only 25 genes were upregulated across all immune cells in the WS group, and these were associated with mitochondrial electron transport, amino acid metabolism, and cytoskeletal organization (Figure 2D). This minimal upregulation suggests a limited and potentially insufficient compensatory transcriptional response to WS.

**Figure 2.**
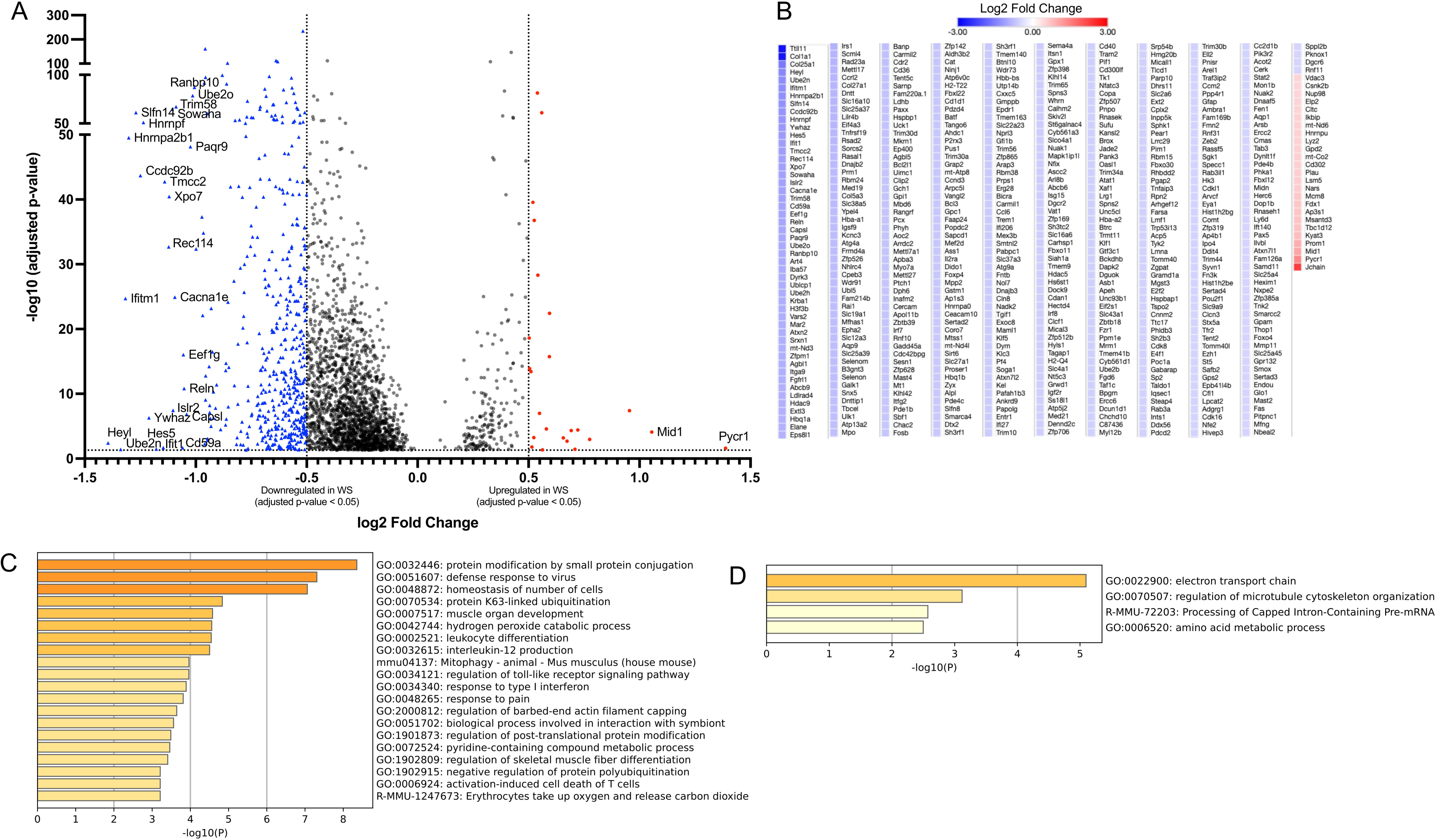
Global transcriptional alterations across bone marrow immune cells in OVX mice after WS exposure. (A) Volcano plot of differentially expressed genes (DEGs) for all bone marrow immune cells, comparing wood smoke (WS)-exposed vs. filtered air (FA)-exposed ovariectomized (OVX) mice (n = 3 per group). Significantly upregulated (red circles) and downregulated (blue triangles) genes are highlighted, respectively (FDR-adjusted p < 0.05 with |logLFC| > 0.5). (B) Heatmap showing the expression patterns of all significant DEGs (FDR-adjusted *p* < 0.05 with |logLFC| > 0.5) between the WS and FA groups. Relative upregulation and downregulation of gene expression are indicated by the color scale. (C, D) Bar graphs showing enriched biological pathways among genes (C) downregulated or (D) upregulated across the immune cell compartment in WS vs. FA. In both panels, the color intensity of each bar corresponds to its *p*-value significance, with darker shades indicating more significant pathways. See also Supplementary Figure S7 for similar transcriptional suppression patterns in specific innate immune cell subsets (neutrophils, monocytes, dendritic cells).

As a likely consequence of this broad transcriptional disruption to immune homeostasis, we also observed subtle but statistically significant shifts in the proportions of specific immune populations (Figure 3A-C). To test these compositional changes while accounting for mouse-to-mouse variability, we used a mixed-effects logistic regression model comparing WS versus FA for each immune cell type. This model identified a significant increase in memory CD8L T cells in WS-exposed OVX mice (odds ratio = 1.23, p < 0.0001; mean proportions 5.13% vs. 6.26%), along with significant decreases in ISG-expressing (interferon-stimulated gene–expressing) immune cells (odds ratio = 0.63, p < 0.0001; 5.82% vs. 3.76%) and granulocytes (odds ratio = 0.60, p < 0.0001; 0.76% vs. 0.45%) compared with FA controls. Smaller but statistically significant increases were also observed in myeloid dendritic cells (odds ratio = 1.25, p = 0.0173; 1.82% vs. 2.26%) and CD8L NKT-like cells (odds ratio = 1.27, p = 0.0012; 2.24% vs. 2.83%). Taken together with the global transcriptional suppression, these compositional changes suggest that WS exposure alters myeloid progenitor function and reshapes the bone marrow immune landscape in ovariectomized mice, consistent with emerging concepts of innate immune memory (Kaufmann et al. 2018; Ochando et al. 2023). The comprehensive results of this model, including violin plots of sample fractions and a forest plot of odds ratios, are presented in Supplementary Figure S3. The raw cell counts per individual mouse and the full statistical output table are provided in Supplementary Table S3 and S4, respectively.

**Figure 3.**
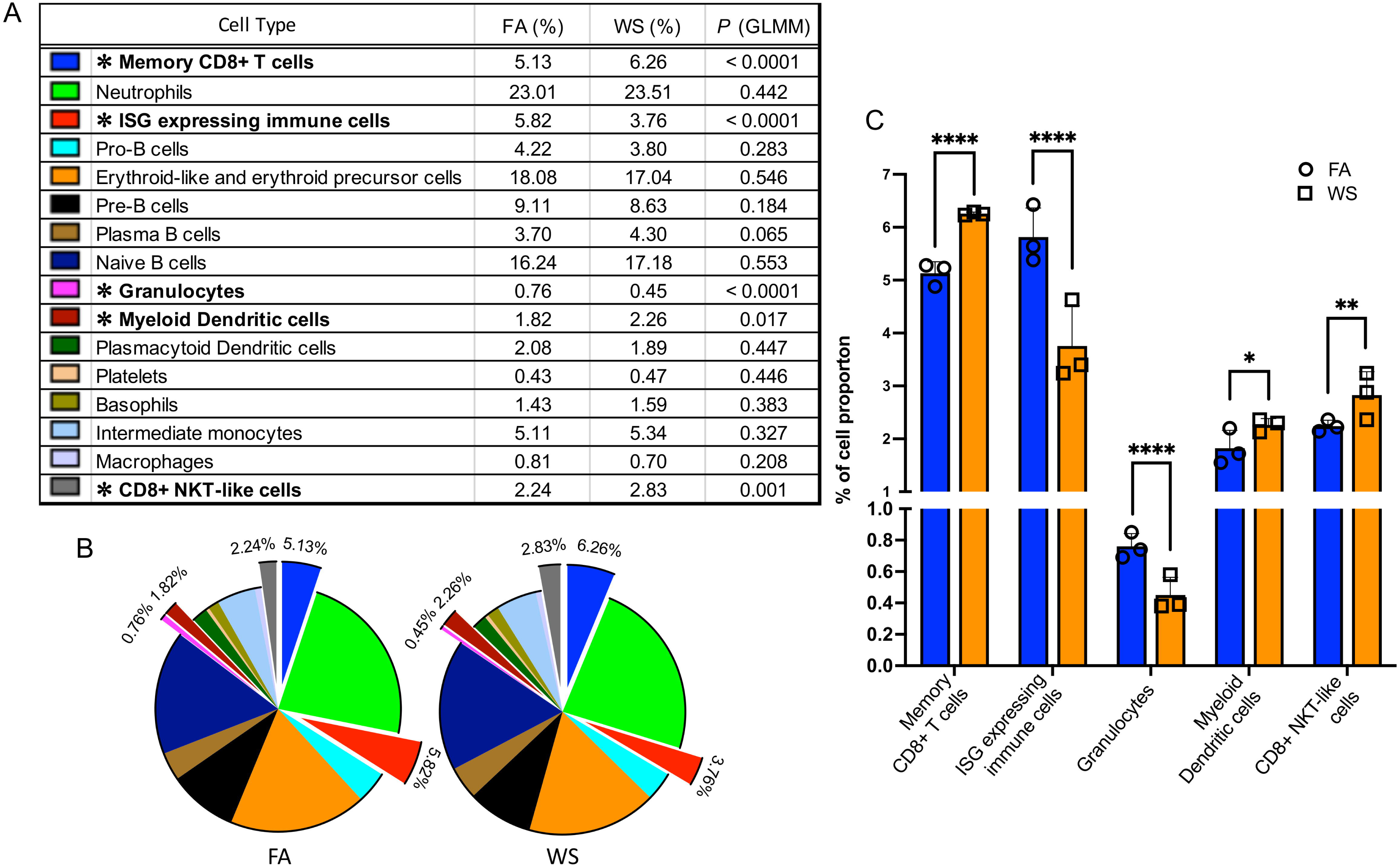
Wood smoke exposure alters bone marrow immune cell population proportions in ovariectomized (OVX) mice. (A) Table comparing the proportions of each bone marrow immune cell type in filtered air (FA) vs. wood smoke (WS) groups (OVX mice, n = 3 per group). *P* values and asterisks indicate significance from a mixed-effects logistic regression model comparing WS versus FA while accounting for inter-animal variability. (B) Pie charts showing the relative composition of immune cell populations under FA and WS conditions. Exploded segments correspond to cell types with significant WS-induced compositional changes in the mixed-effects model (asterisked in A). (C) Bar and scatter plots highlighting immune cell types with significant differences in proportions between FA and WS: memory CD8L T cells, granulocytes, ISG-expressing (interferon-stimulated gene-expressing) immune cells, myeloid dendritic cells, and CD8L NKT-like cells. Symbols represent per-mouse proportions and bars indicate group means ± SD (n = 3 per group). Statistical significance (asterisks) is based on the mixed-effects logistic regression model: *p < 0.05; **p < 0.01; ***p < 0.001; ****p < 0.0001 (Supplementary Tables S3-S4).

### Suppressed immune activation pathways in altered cell subsets

In memory CD8L T cells, we identified a total of 41 significant differentially expressed genes (DEGs) (adjusted *p* < 0.05, |logLFC| > 0.5) within the OVX context (Figure 4A). Functional enrichment analysis of these genes revealed that downregulated DEGs were associated with key immune functions, including leukocyte activation, while upregulated DEGs were linked to cellular response to stress. We visualized the effect sizes (median logLFC and 95% CI) for the 18 representative genes from these top pathways, confirming the direction and confidence of these changes (Figure 4C, 4E). A heatmap of this 18-gene signature demonstrated a clear separation between the FA and WS groups (Figure 4F). This finding was confirmed at the animal-replicate level (Supplementary Figure S5A). We performed pseudobulk pathway analysis (using a relaxed *p* < 0.05 cutoff appropriate for n=3 replicates) and successfully recapitulated the significant suppression of the exact same biological pathways, including leukocyte migration (-Log10*p*=9.57) and leukocyte activation (-Log10*p*=7.48). This analysis also recapitulated the activation of stress-response pathways (e.g., regulation of DNA metabolic process, DNA repair). Furthermore, unbiased GSEA (Supplementary Figure S6A) robustly validated this conclusion. GSEA revealed a broad, statistically significant suppression (NES < -0.5, FDR < 0.001) of the entire immune response and leukocyte migration pathways, alongside a strong activation (NES > 0.5, FDR < 0.001) of DNA repair and RNA processing stress-response pathways.

**Figure 4.**
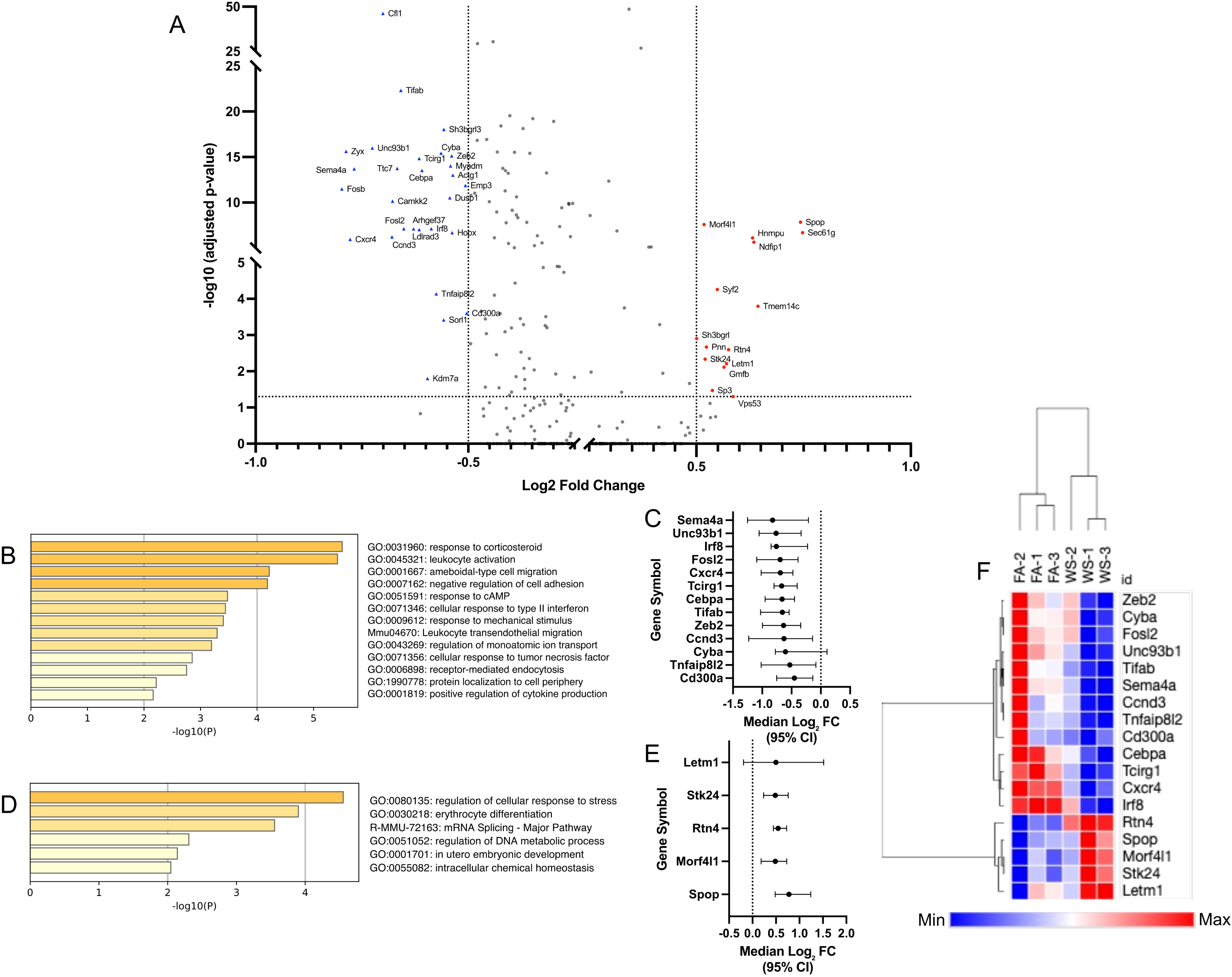
Wood smoke (WS)–induced transcriptional changes in memory CD8L T cells derived from OVX mice bone marrow. (A) Volcano plot of differentially expressed genes (DEGs) in memory CD8L T cells from ovariectomized (OVX) mice (WS vs. FA conditions, n = 3 mice per group). Significantly upregulated (red circles) and downregulated (blue triangles) genes are highlighted. Key representative genes are labeled by name. Significance cutoffs were set at FDR-adjusted p < 0.05 and |logLFC| > 0.5, which identified a total of 41 significant DEGs. (B, D) Functional enrichment analysis of the 41 significant DEGs from (A). Panel (B) shows the top pathways for downregulated genes, while panel (D) shows pathways for upregulated genes. In both panels, the color intensity of each bar corresponds to its p-value significance, with darker shades indicating more significant pathways. (C, E) Forest plot showing the effect sizes (median logLFC and 95% CI) for the 18 key representative genes selected from the top biological pathways identified in (B) and (D) (e.g., leukocyte activation and cellular response to stress). (F) Heatmap of the 18 core pathway genes from (C) and (E), generated using row-based z-scores. Unsupervised clustering of samples illustrates clear separation of FA and WS within this cohort (n=3 per group).

In parallel, ISG-expressing immune cells displayed an even stronger transcriptional response to WS exposure in the OVX context. We identified 121 significant DEGs, which were enriched for the suppression of protein modification (e.g., ubiquitination) pathways and the potent upregulation of chromatin remodeling and cell cycle pathways (Figure 5B, 5D). Effect sizes (Figure 5C, 5E) and heatmap analysis (Figure 5F) confirmed this robust signature. This finding was also confirmed at the animal-replicate level (Supplementary Figure S5B). Pseudobulk pathway analysis recapitulated the suppression of modification-dependent protein catabolism (-Log10*p*= 8.86) and response to virus, and the strong activation of Cell Cycle (-Log10*p*=6.1) and chromatin remodeling pathways. Finally, GSEA provided a robust, orthogonal validation, confirming the suppression of cell-cell signaling (NES < 0, FDR < 0.001) and the overwhelming activation of chromosome organization (NES: +6.67, FDR 0.0) (Supplementary Figure S6B).

**Figure 5.**
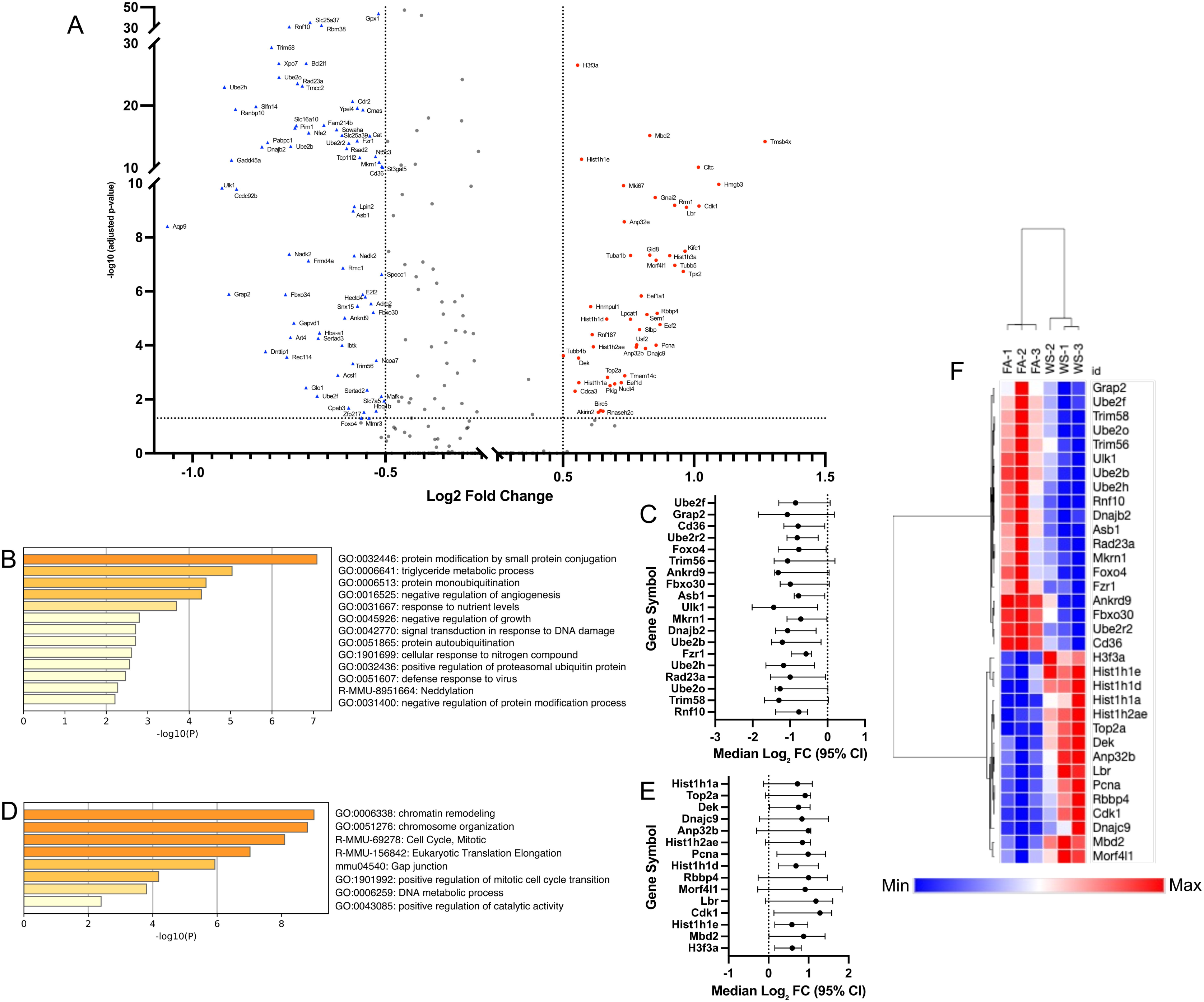
Wood smoke (WS)–induced transcriptional dysregulation in ISG-expressing immune cells from OVX mouse bone marrow. (A) Volcano plot of differentially expressed genes (DEGs) in ISG-expressing immune cells from ovariectomized (OVX) mice (WS vs. FA conditions, n = 3 mice per group). Significance cutoffs (FDR-adjusted p < 0.05, |logLFC| > 0.5) are indicated by dashed lines. Key representative genes are labeled by name. (B, D) Functional enrichment analysis of the 121 significant DEGs identified in (A). (B) shows the top-ranked biological pathways for down-regulated genes, while (D) shows the top-ranked pathways for up-regulated genes. Color intensity corresponds to p-value significance. (C, E) Forest plots showing the effect sizes (median logLFC and 95% CI) for the core set of genes selected from the top-ranked pathways in (B) and (D). (C) displays genes from the ‘protein modification by small protein conjugation’ pathway (19 genes). (E) displays genes from the ‘chromatin remodeling’ related pathways (15 genes). (F) Heatmap of the 34 core pathway genes from (C) and (E), generated using row-based z-scores. Hierarchical clustering of samples (columns) demonstrates that this core biological signature separates all FA (n=3) from WS (n=3) replicates.

For complete data transparency and to support the curated validation analyses, the full uncurated data are provided. Forest plots for all remaining significant DEGs not included in the main figures are provided in Supplementary Figure S4. The full Top 20 pathway lists from the Pseudobulk validation (Supplementary Figure S5) are provided in Supplementary Table S6. Furthermore, the full lists of all significant pathways (FDR < 0.01) from the GSEA validation (Supplementary Figure S6) are provided in Supplementary Table S7.

Finally, Broader impacts on other innate immune cells, including neutrophils, monocytes, and myeloid dendritic cells showed similar transcriptional suppression in pathways governing neutrophil degranulation, cytokine signaling, and type I interferon responses (Supplementary Figure S7).

### Ovarian hormone deficiency sensitizes BMDMs to WS-induced metabolic reprogramming

To determine if this transcriptional reprogramming in myeloid precursors alters macrophage function, we next profiled the metabolic activity of BMDMs from all four experimental groups. Successful macrophage differentiation was validated by flow cytometry, with the great majority of harvested adherent cells coLexpressing CD11b and F4/80 (SupplementaryLFigureLS8). Under baseline (FA) conditions, BMDMs from OVX mice showed reduced mitochondrial and glycolytic function compared to Sham controls (Figure 6B-D, G-J). Strikingly, WS exposure in OVX mice caused what is believed to be hypersensitive compensatory response, bringing both mitochondrial respiration and glycolysis back to the levels seen in Sham mice. This effect was entirely dependent on hormonal status, as WS had no impact on the metabolism of macrophages from Sham mice.

**Figure 6.**
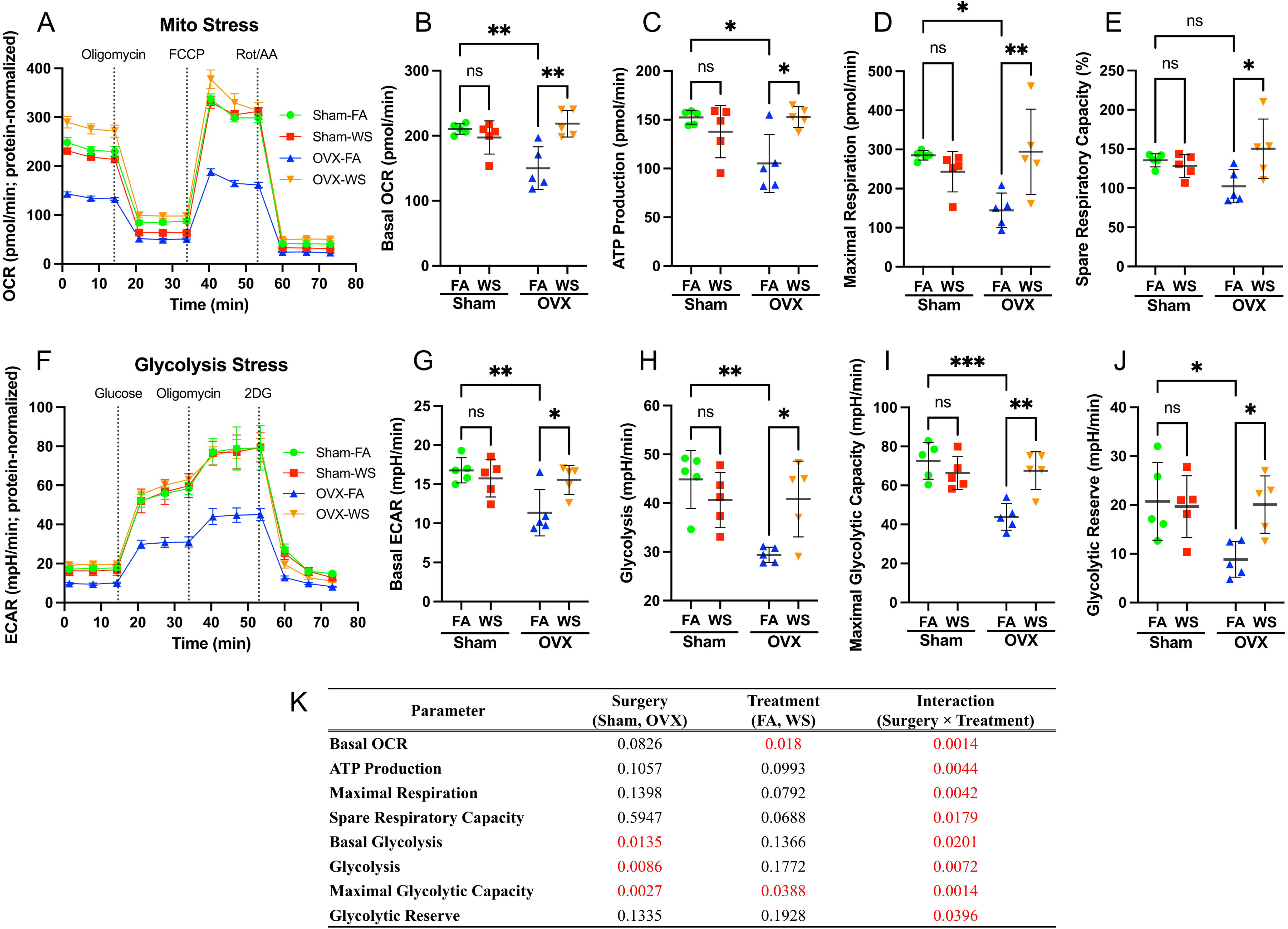
Ovarian hormone deficiency and WS exposure interact to disrupt mitochondrial and glycolytic metabolism in BMDMs. (A) Real-time oxygen consumption rate (OCR) in bone marrow–derived macrophages (BMDMs), measured with the Seahorse XF Mito Stress test. The curve shows mean OCR ± SD over time for BMDMs from Sham and ovariectomized (OVX) mice exposed to filtered air (FA) or wood smoke (WS), with sequential injections of oligomycin, FCCP, and rotenone/antimycin A (mitochondrial stressors) indicated. (B–E) Quantification of OCR parameters derived from (A): (B) basal respiration, (C) maximal respiration, (D) ATP production rate, and (E) spare respiratory capacity. WS exposure effects are compared between Sham and OVX BMDMs for each parameter. (F) Real-time extracellular acidification rate (ECAR) in BMDMs, measured with the Seahorse XF Glycolysis Stress test. The kinetic profile (mean ± SD) is shown for Sham vs. OVX BMDMs under FA or WS exposure, with injections of glucose, oligomycin, and 2-deoxy-D-glucose (2-DG) to probe glycolytic function. (G–J) Quantification of ECAR parameters from (F): (G) basal glycolysis, (H) glycolysis, (I) maximal glycolytic capacity, and (J) glycolytic reserve. Data shown are protein-normalized OCR/ECAR (native units, pmol/min and mpH/min) from wells seeded equally at 80,000 cells. These bar graphs compare metabolic parameters between FA and WS in Sham and OVX groups. Data are presented as scatter plots showing individual data points, with bars representing the mean ± SD (n = 5 mice per group, each data point represents the average of six technical replicates per mouse). Two-way ANOVA with Tukey’s multiple comparisons test was used to analyze the effects of treatment (FA vs. WS) in each group. Significance is indicated as \*\*\**p* < 0.001; \*\**p* < 0.01; \**p* < 0.05; ns = not significant. (K) The main effects of surgery and treatment, and their interaction, were analyzed by two-way ANOVA, followed by Tukey’s multiple comparisons test between all groups on each parameter (from panels B-E, G-J). Significant interaction effects (*p* < 0.05) were observed for all measured metabolic parameters, indicating that the metabolic response to WS exposure is fundamentally dependent on ovarian hormone status.

A two-way ANOVA confirmed a significant Surgery × Treatment interaction for all metabolic parameters (Figure 6K), demonstrating that WS dramatically reprograms macrophage metabolism, but only within the context of ovarian hormone deficiency. For the purposes of this study, we use the term "reprogramming" specifically to denote this functional shift away from a canonical cellular state, rather than to imply a permanent, irreversible fate change. These pronounced metabolic enhancements, when considered alongside the broad transcriptional suppression of immune activation pathways in their progenitor niche (Figure 2C), suggest that WS exposure in ovarian hormone-deficient mice reprograms metabolic pathways independently of classical immunoregulatory gene expression, consistent with the concept of immune–metabolic decoupling.

### Wood smoke exposure selectively curtails M2-associated gene expression in macrophages from ovarian hormone-deficient mice

Crucially, the macrophage response to WS depended on ovarian hormone status. This conclusion is supported by a significant Surgery × Treatment interaction for most M2-associated genes in a linear mixed-effects model (LMM), which accounts for inter-mouse variability (Figure 7H). Post-hoc tests revealed that WS exposure led to a potent and significant downregulation of key M2-associated genes, including *Arg1*, *Mrc1*, *Pparg*, *Tgfb1*, and *Ccl2*, but only in BMDMs derived from OVX mice (Figure 7A-E). In stark contrast, this transcriptional suppressive effect was absent in Sham mice. Consistent with the downregulation of M2-associated genes in differentiated macrophages (Figure 7), our global scRNA-seq analysis across all bone marrow immune cells revealed a significant suppression of the key upstream M2-polarizing transcription factors, including *Stat6*, *Klf4*, and *Irf4*, as well as the PPARγ-regulated scavenger receptor *Cd36*, in WS-exposed OVX mice compared to FA controls (Supplementary Table S5).

**Figure 7.**
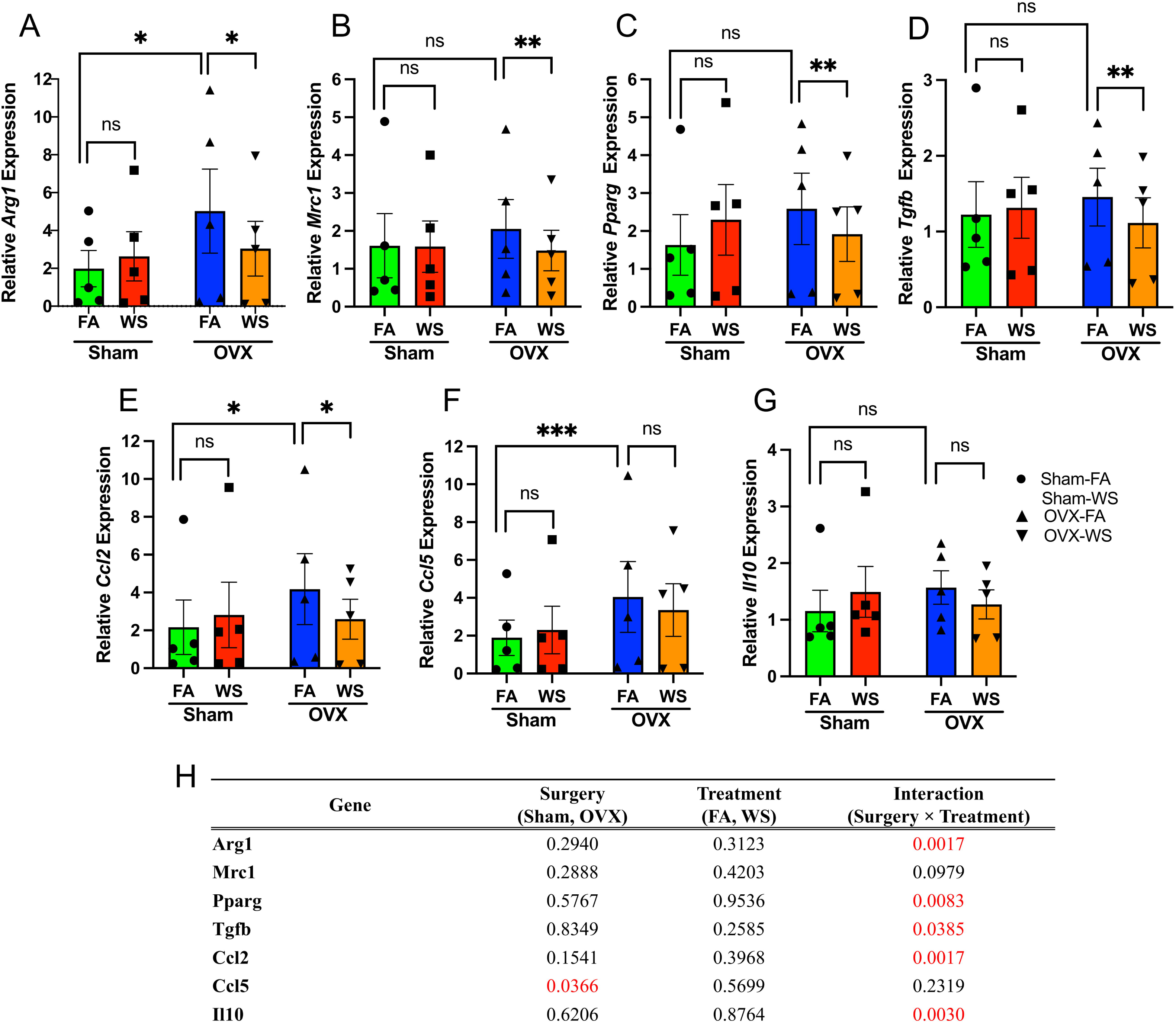
Ovarian hormone deficiency and WS exposure modulate M2-associated gene expression in BMDMs. (A–G) Relative mRNA expression of M2-associated genes in bone marrow–derived macrophages (BMDMs) from Sham and ovariectomized (OVX) mice exposed to filtered air (FA) or wood smoke (WS). BMDMs were differentiated from bone marrow of each group and gene expression was measured by quantitative RT-PCR. Panels show expression of (A) *Arg1*, (B) *Mrc1*, (C) *Pparg*, (D) *Tgfb1*, (E) *Ccl2*, (F) *Ccl5*, and (G) *Il10*. For visualization, gene expression levels were normalized to 18S rRNA and relative quantification was calculated using the 2−ΔΔCt method, with the mean ΔCt of the Sham-FA group serving as the calibrator for all samples. Bars represent mean ± SEM for each group, and each data point represents an individual mouse (n = 5 biological replicates per group, with 3 technical replicate measurements averaged per mouse). (H) Summary of *p*-values for the main effects and their interaction from the two-way mixed-effects model. Statistical significance for all panels was determined using a linear mixed-effects model (two fixed factors: Surgery, Treatment; random intercept: Biological replicate) on the ΔCt values (prior to 2−ΔΔCt transformation) to appropriately account for inter-replicate variability. ‘Surgery’ and ‘Treatment’ were modeled as fixed effects, and ‘Biological Replicate’ as a random effect. Asterisks denote significant differences from post-hoc multiple comparisons tests. *\*\*\*p* < 0.001; *\*\*p* < 0.01; *\*p* < 0.05; ns = not significant. See also Supplementary Figure S10 for the response patterns of each individual biological replicate. See also Supplementary Figure S11 for data showing that expression of representative M1-associated genes (e.g., *Cxcl1*, *Il1b*, *Tnfa*) was not significantly altered by WS under these conditions.

Interestingly, this differential response to WS appears to be layered upon a pre-existing hormonal influence. Under baseline (FA) conditions, OVX BMDMs showed lower mitochondrial and glycolytic capacity than Sham (Figure 6B–E, G–J), whereas several M2-linked transcripts (e.g., *Arg1, Ccl2, Ccl5*) and proteins were higher at baseline in OVX-FA versus Sham-FA (Figure 7A, E–F; Figure 8B–C). Together, these findings indicate that OVX establishes a distinct baseline state characterized by reduced bioenergetics and a partial M2 skew, independent of WS, which likely conditions the subsequent interaction-type response to exposure.

**Figure 8.**
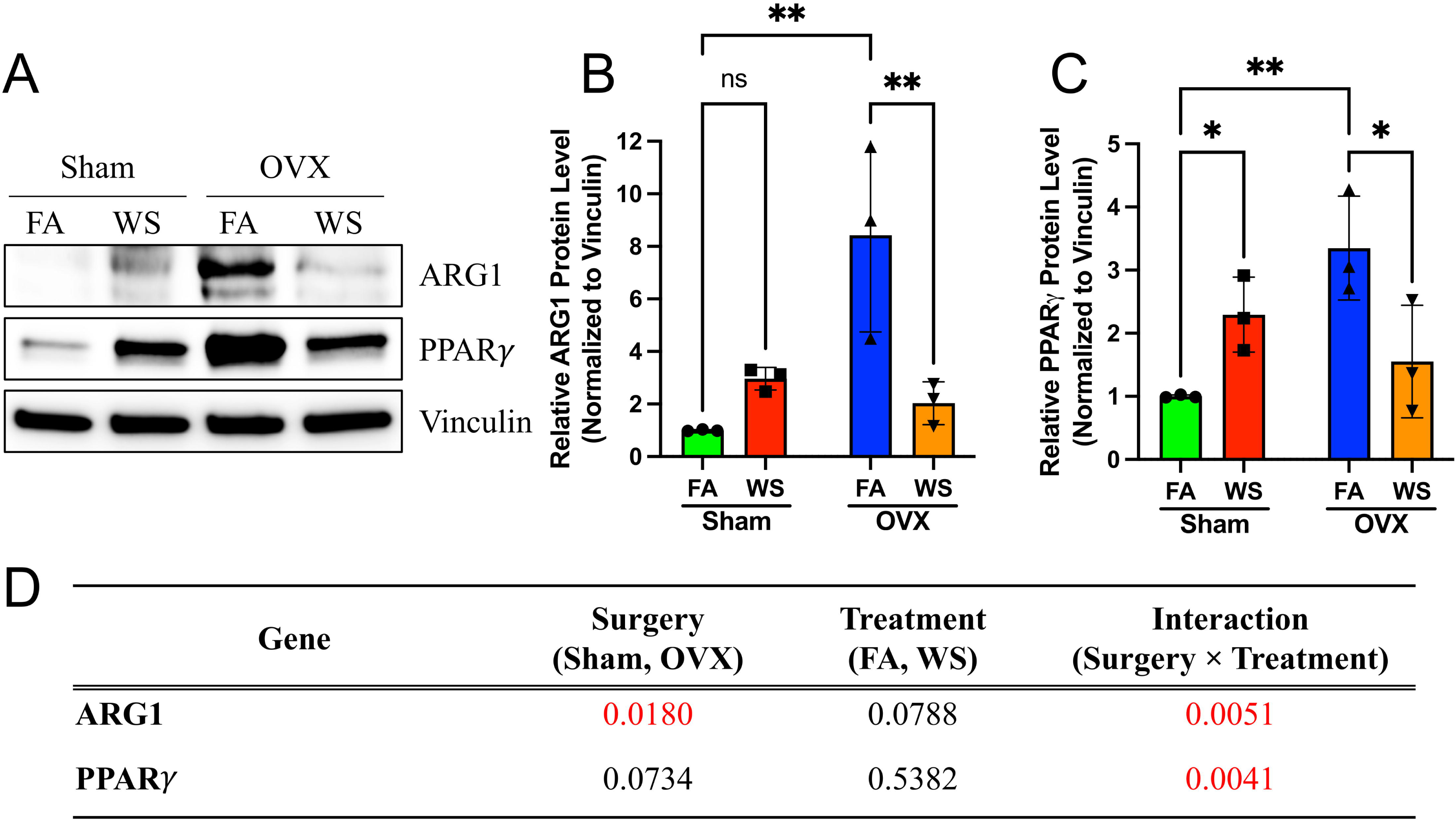
Protein level validation confirms downregulation of ARG1 and PPARγ in OVX-WS BMDMs. (A) Representative Western blot images for ARG1, PPARγ, and Vinculin (loading control) in bone marrow–derived macrophage (BMDM) lysates from Sham and ovariectomized (OVX) mice exposed to filtered air (FA) or wood smoke (WS) (n=3 biological replicates shown). (B) Quantification of relative ARG1 protein levels normalized to Vinculin. (C) Quantification of relative PPARγ protein levels normalized to Vinculin. Data in B and C are presented as scatter plots showing individual biological replicates (n=3), with bars representing mean ± SD. Statistical significance was determined by Two-way ANOVA followed by Tukey’s multiple comparisons test (*p < 0.05, **p < 0.01 compared to OVX-FA, mark specific comparisons based on post-hoc results). (D) Summary table of Two-way ANOVA results showing the P values for the main effects of Surgery, Treatment, and their Interaction on ARG1 and PPARγ protein levels. The significant interaction effects confirm the specific downregulation in the OVX-WS group.

The selective suppression of the M2-associated transcriptional signature did not coincide with a shift toward a pro-inflammatory state, as the expression of representative M1-associated genes (*Cxcl1*, *Il1b*, *Tnfa*) was not significantly altered by WS in either group (Supplementary Figure S11). For clarity, we refer to these gene signatures as M1- and M2-associated genes, recognizing that they reflect transcriptional programs rather than polarization induced by canonical stimuli.

Consistent with the mRNA data, we observed a significant reduction in the protein levels of the key M2 effector enzyme ARG1 and the master regulator PPARγ specifically in macrophages derived from WS-exposed OVX mice (Figure 8A-C). Two-way ANOVA confirmed highly significant Surgery × Treatment interaction effects for both ARG1 (Interaction p = 0.0051) and PPARγ (Interaction p = 0.0041) (Figure 8D), reinforcing that the suppression of the M2-associated program occurs at the protein level and is dependent on ovarian hormone status. These findings support a functionally dysregulated, immune–metabolically decoupled phenotype in OVX under WS, rather than classical M1 induction.

## Discussion

This study provides a mechanistic framework for the systemic immunotoxicity of inhaled pollutants, demonstrating that acute WS exposure, particularly in ovarian hormone deficiency, is a potent suppressor of the bone marrow immune niche. The most striking initial finding in OVX mice was the broad downregulation of immune-related gene programs across hematopoietic lineages (Figure 2), a finding that aligns with and extends previous reports on air pollutant immunotoxicity (Bhattarai et al. 2024; Gangwar et al. 2020). This profound suppression of type I interferon response pathways underscores the severe immunosuppressive consequences of WS exposure. Given the critical role of interferon-stimulated genes in antiviral defense, their downregulation may provide a mechanistic basis for the increased incidence of respiratory infections observed following wildfire smoke exposure (Brocke et al. 2022; Frumento and Țãlu 2024). Furthermore, this immune disruption was evident in the significant compositional shifts within the bone marrow niche. Statistical modeling of cell-type proportions revealed an increase in memory CD8L T cells, reductions in granulocyte and ISG-expressing cell populations, and significant increases in myeloid dendritic cells and CD8+ NKT-like cells (Figure 3; Supplementary Figure S3B; Supplementary Table S4). These widespread compositional changes, coupled with suppressed degranulation pathways (Supplementary Figure S7), suggest impaired myeloid differentiation and compromised inflammatory responses, which could heighten the risk of infections (Frumento and Țãlu 2024; Migliaccio et al. 2013).

While memory CD8L T cells increased in number (Figure 3), their transcriptional profile indicated functional impairment within the OVX context. Our cell-level analysis identified the suppression of key immune activation pathways (Figure 4). This finding was comprehensively validated by two independent, orthogonal methods. First, animal-level pseudobulk pathway analysis confirmed the suppression of the exact same leukocyte activation and migration pathways (Supplementary Figure S5A). Second, unbiased GSEA robustly validated the broad suppression of the immune response pathway (Supplementary Figure S6A). Notably, this validation effort also highlighted a specific mechanistic disruption: the pseudobulk analysis identified the upregulation of the chemokine *Cxcl12*, which coincides with the suppression of its receptor, *Cxcr4*, observed in our single-cell analysis. This pattern suggests a disruption of the CXCL12-CXCR4 signaling axis, a critical pathway for T-cell trafficking and retention in the bone marrow (Zhao et al. 2012), and such disruption could contribute to impaired immune surveillance and increased risk of immune-related pathology (Mantovani et al. 2008; Scharf et al. 2020).

This transcriptional dysregulation was also evident in ISG-expressing cells, key mediators of tumor surveillance. The suppression of protein modification (ubiquitination) pathways and the profound activation of chromatin remodeling and cell cycle pathways were identified at the cell-level (Figure 5) and validated at both the animal-replicate pseudobulk (Supplementary Figure S5) and whole-transcriptome (GSEA) levels (Supplementary Figure S6). This comprehensive validation of suppressed antiviral and protein-modification pathways, alongside induced genomic stress, is consistent with mechanisms that have been implicated in increased cancer susceptibility, in line with epidemiologic data linking WS exposure to hematopoietic malignancies (De Guzman and Schiller 2025; Hohtari et al. 2019; Maurer et al. 2025; Navarro et al. 2019). Finally, the disruption of pathways related to bone remodeling (Supplementary Figure S5A) and mitochondrial homeostasis (Supplementary Figure S6A) aligns with clinical and experimental data linking air pollution to reduced bone density and hematopoietic stem cell dysfunction (Bhattarai et al. 2024; Prada et al. 2023).

A key strength is the compelling evidence for the *in vivo* priming of progenitor cells, as macrophages differentiated *ex vivo* retained a memory of the *in vivo* exposure condition. This suggests that WS exposure induces lasting alterations within the bone marrow myeloid compartment. This interpretation is further supported by the significant interaction between surgical status (Sham vs OVX) and exposure (FA vs WS) in functional BMDM assays, which rigorously included Sham controls, confirming that the observed phenotype is hormonally dependent. These alterations could reflect either cell-intrinsic imprinting or WS-induced shifts in monocyte precursor subtypes. However, the persistence of pronounced metabolic and transcriptional differences after normalization for cell number lends strong support to the hypothesis of a cell-intrinsic reprogramming, though the duration of this impact remains unknown.

These findings fit within the trained immunity framework whereby environmental stimuli remodel bone marrow progenitors, altering downstream immune function (Kaufmann et al. 2018; Liao et al. 2025). Similarly, environmental toxicants have been shown to epigenetically modify hematopoietic stem and progenitor cells (HSPCs), resulting in lineage-specific biases or dysregulated inflammatory responses in differentiated cells (Bhattarai et al. 2024; Scharf et al. 2020). Our findings extend these principles to air pollution, demonstrating that WS exposure, especially in an ovarian hormone-deficient state, exerts a lasting influence on immune cell development and function via the bone marrow niche.

Our functional analyses revealed that the macrophage response to WS was fundamentally dictated by ovarian hormone status. BMDMs from WS-exposed OVX mice exhibited a profound metabolic reprogramming that reversed their otherwise suppressed metabolic state, an effect absent in Sham mice (Figure 6). Paradoxically, this metabolic activation in OVX-derived macrophages was accompanied by a selective dismantling of their M2-associated gene program (e.g., *Arg1*, *Mrc1, Pparg*, *Tgfb1, Ccl2*), while M1-associated genes remained unchanged (Figure 7, Supplementary Figure S11). This striking divergence in response, metabolic activation alongside selective immune gene suppression, occurring only under conditions of ovarian hormone deficiency, provides a clear demonstration of immune–metabolic decoupling. We therefore operationally define this phenotype as heightened mitochondrial/glycolytic flux concomitant with suppression of M2-associated transcription in the absence of M1 induction. Importantly, the OVX baseline itself, a complex phenotype characterized by reduced bioenergetics alongside a partial M2 skew under FA conditions, likely conditions the subsequent WS phenotype. The pre-existing metabolic suppression might render these cells less flexible or poised for a more drastic compensatory response upon encountering a metabolic challenge like WS. Consistent with this, WS elicited a surgery-dependent interaction: immune–metabolic decoupling emerged in OVX but not in Sham (Figure 6–8). This framework explains why estrogen-replete Sham remained largely buffered, whereas OVX, already existing in an altered metabolic and transcriptional state, manifested the combined metabolic activation and selective M2 suppression upon WS exposure.

Mechanistically, PPARγ attenuation appears central (Bouhlel et al. 2007; Odegaard et al. 2007). In macrophages from WS-exposed OVX mice, *Pparg* mRNA and PPARγ protein were reduced (Fig. 7, Fig. 8C), consistent with decreases in *Arg1/Mrc*1 mRNA (Fig. 7A–B) and ARG1 protein (Fig. 8B) levels. The concept that environmental toxicants can suppress PPARγ signaling finds parallels in other contexts, such as arsenite-mediated downregulation of *Pparg* (Li et al. 2022). Given PPARγ’s role in alternative activation and in mitochondrial biogenesis/fatty-acid oxidation (Chawla 2010; Vats et al. 2006), the concurrent rise in mitochondrial/glycolytic flux despite lower PPARγ suggests a hormone-dependent metabolic reconfiguration rather than canonical M2 programming. The established role of estrogen in modulating alternative activation (Markle and Fish 2014; Ray et al. 2024; Townsend et al. 2012) further supports this interpretation. Taken together, we propose a working model in which WS exposure in a hormoneLdeficient state could attenuate PPARLγ–linked M2 programs and bias cellular energetics toward compensatory, partially uncoordinated pathways, potentially involving mTOR/HIFL1α–linked glycolytic and epigenetic remodeling circuits (Liu et al. 2017; Muthumalage and Rahman 2023; Tannahill et al. 2013). Because we did not manipulate PPAR-γ or mTOR/HIFL1α directly, these pathways remain provisional and will require targeted perturbation in future work. Even so, the present correlative evidence supports a working model in which endocrine context conditions the metabolic plasticity of macrophages, yielding the observed decoupled phenotype.

Given the critical role of estrogen, a plausible alternative hypothesis is that components of WS itself might exert estrogenic effects, as has been observed for other environmental toxicants (Lascombe et al. 2000; Peng et al. 2023). Notably, our data do not support an additive estrogenic effect from WS exposure. Such an effect would predict that WS exposure would enhance the M2-program in estrogen-replete Sham mice; instead, we observed no such additive effect in this group. Our data strongly support a hormone-status–dependent interaction: in estrogen-replete Sham mice, the PPARγ-linked alternative-activation axis is largely buffered, whereas in OVX marrow, WS preferentially depresses this axis, yielding the observed decoupling (heightened mitochondrial/glycolytic flux with selective M2 suppression) (Gonzalez et al. 2023; Viola et al. 2019).

This immune–metabolic decoupling is the co-occurrence of metabolic activation (Figure 6) and selective M2-associated transcriptional and protein downregulation (Figure 7, 8), all occurring in the absence of M1 induction. This finding challenges the canonical tight coupling between metabolism and polarization (Liu et al. 2017; Viola et al. 2019) and aligns with emerging reports of context-dependent decoupling, notably resembling patterns in tumor-associated macrophages where robust metabolic activity coexists with immunosuppressive functions (Gonzalez et al. 2023; Liu et al. 2023). It is analogous to findings in fibrotic lung injury models, where distinct macrophage subsets exhibit divergent metabolic and transcriptional states (Moos et al. 2024). Notably, similar immune–metabolic decoupling and potential involvement of mTOR signaling have been reported in other inhalation models; for example, flavored e-cigarette aerosols have been shown to induce metabolic rewiring of T cells via PI3K/Akt/mTOR signaling, accompanied by suppression of immune-related gene expression (Muthumalage and Rahman 2023). Furthermore, this phenotype can be discussed within the broader conceptual framework of innate immune memory, which includes trained tolerance, an arm where prior stimulation leads to a dampened inflammatory response (Divangahi et al. 2021). It is therefore plausible that in the absence of the regulatory influence of ovarian hormone, WS exposure does not induce classical pro-inflammatory training but instead primes myeloid progenitors toward a metabolically active but immunologically non-canonical state (Saaoud et al. 2023), a phenotype linked to detrimental outcomes like impaired pathogen clearance and tumor progression (Migliaccio et al. 2013; Wynn and Vannella 2016). This bone marrow priming could be considered an early key event in an adverse outcome pathway leading to long-term immune dysfunction. Importantly, these findings elevate the role of bone marrow progenitors from passive precursors to active integrators of environmental and endocrine signals. This has broad implications for environmental immunotoxicology and public health, raising the possibility that transient exposures during sensitive physiological windows, such as the postmenopausal period, may contribute to long-term alterations in innate immune readiness and inflammatory tone via bone marrow programming.

Such lasting alterations in bone marrow progenitors likely involve systemic signals originating from the lung, such as circulating inflammatory mediators, consistent with our prior lipidomic findings (Wardhani et al. 2024a) and as reviewed in (Scharf et al. 2020), extracellular vesicles carrying molecular cargo (Lopez et al. 2022), or neuro-endocrine signaling pathways that modulate the hematopoietic niche (Kodavanti et al. 2023). Furthermore, the direct translocation of ultrafine/soluble constituents remains plausible (Bhattarai et al. 2024), given our WS aerosol MMD of 0.13 µm (Supplementary Table S1).

This study focuses on an acute murine model and may not capture the full spectrum of chronic human exposures. We also inferred cell-population shifts primarily from transcriptomic data rather than cytometric validation; while patterns were consistent across replicates, complementary flow cytometry in future work would further strengthen these findings. Our data provide a single time-point snapshot; although differences persisted through seven days of *ex vivo* culture, the long-term stability of this imprint *in vivo* remains to be determined. We cannot yet distinguish whether the phenotype reflects epigenetic imprinting within specific progenitors or shifts in precursor subtypes; targeted sorting of defined progenitors will be required. A key methodological limitation is the absence of a Sham-operated cohort in the scRNA-seq analysis, which prevents direct verification of the interaction between surgical status and exposure at single-cell resolution. Accordingly, we restrict single-cell interpretations to the OVX context and rely on orthogonal functional assays, including both Sham and OVX cohorts, to support interaction effects (Figure 6, 7, and 8); future studies will extend single-cell profiling to Sham cohorts to directly test this interaction at cellular resolution. We acknowledge that we did not stage the estrous cycle in Sham females; however, the impact of this limitation is substantially mitigated by evidence confirming the intended robust endocrine contrast: this was verified by OVX-induced weight gain in the present cohort (Supplementary Figure S1; a key physiological validation of estrogen deficiency) and prior confirmation of regular cycling in ovary-intact females in our prior work (Wardhani et al. 2024b). In the downstream macrophage functional assays, although bead/FACS re-enrichment can increase uniformity, the required sorting and re-plating introduce viability/activation and re-adhesion variability that can confound Seahorse metabolic readouts; therefore, we used an identical differentiation/plating workflow across groups and document maturation-dependent increases in F4/80 as procedural quality control (Supplementary Figure S8).

While our results define a distinct immune-metabolic state in macrophages, the functional consequences of this phenotype *in vivo* require further investigation. Intriguingly, this immune-metabolically uncoupled state, defined by the suppression of the M2-associated program without M1 polarization, presents a central paradox given our own prior work demonstrating that OVX mice exposed to wood smoke exhibit systemic hyper-inflammation (Wardhani et al. 2024a; Wardhani et al. 2024b). This raises the critical mechanistic question of how these non-canonically programmed macrophages drive a pro-inflammatory systemic environment. We propose that this "decoupled" phenotype defines a novel pathogenic state, contributing to systemic pathology not by classical activation, but by failing to resolve inflammation or by engaging in dysregulated crosstalk with other immune cells (e.g., neutrophils or T cells). Therefore, assessing the functional consequences of this state, such as the response to secondary stimuli (e.g., LPS) or behavior in co-culture systems, represents a crucial next step to delineate this aberrant mechanism and its contribution to systemic pathology.

Our findings demonstrate that ovarian hormone deficiency sensitizes the bone marrow to WS-induced immune disruption, characterized by transcriptional suppression, immune compositional shifts, and uncoupled metabolic reprogramming in macrophages. These results underscore the importance of ovarian hormonal status in modulating systemic responses to air pollution. Indeed, the need to understand the biological and health effects of fire smoke is increasingly highlighted as a critical area of public health (Oh and Zychowski 2025). Our findings contribute to this imperative by providing deeper mechanistic insight into WS-induced immunotoxicity, which is crucial for developing more precise risk assessments for hormonally susceptible populations.

## Data Availability

The raw single-cell RNA sequencing data reported in this paper have been deposited in the Dryad digital repository, under accession number DOI: 10.5061/dryad.bvq83bknp

## Acknowledgments

Dr. Zychowski would like to thank the UNM-INSPIRES Translational Resource Support Core (TRSC) and the Campen Lab for usage of the WS inhalation system (P30 ES032755). Our team would also like to acknowledge the AIM Center at UNM (P20 GM1121176) and the IMSD Program (T32 GM144834).

## Author contributions

Conceptualization, M.O., K.E.Z.; methodology, A.B., and S.Y.; investigation, M.O., E.L, O.E, C.M., A.B., J.M.G, and S.Y.; data curation, M.O., S.Y., and E.L., writing-original draft, M.O.; writing-review & editing, M.O., J.M.G, A.B., and S.Y.; funding acquisition, K.E.Z. and S.Y; resources, K.E.Z.; supervision, K.E.Z

## Supplementary Material

Document S1. Supplementary Table S1-S5 and Supplementary Figures S1–S11

Supplementary Tables S6 and S7 (excel files). These files contain all large-scale transcriptomic data supporting the validation analyses. Supplementary Table S6 provides the pseudobulk input gene list and Metascape pathway results, and Supplementary Table S7 provides the full Gene Set Enrichment Analysis (GSEA) pathway results.

