## Supplementary Materials for "Ovarian hormone deficiency enhances wood smoke-induced immune dysfunction via transcriptomic and metabolic alterations"

**Supplementary Matarials**

**Supplementary Table S1. WS particle-size summary (Aerosizer)**

| Metric | Value |
| --- | --- |
| Mass median diameter (MMD), μm | 0.13 |
| Geometric standard deviation (GSD) | 1.4 |
| Sampling protocol | 100 one-minute samples |

Numeric recap of WS particle-size metrics previously measured with the identical generator settings used here (Wardhani et al. 2024).

**Supplementary Table S2. Particle-phase metals in WS vs FA (ICP-MS)**

| Element (isotope) | P-value | Element (isotope) | P-value |
| --- | --- | --- | --- |
| ²³Na | 0.5373 | ⁷⁵As | 0.4816 |
| ²⁴Mg | 0.4640 | ⁷⁸Se | 0.8548 |
| ²⁷Al | 0.4428 | ¹¹¹Cd | 0.4106 |
| ³⁹K | 0.4862 | ⁴⁴Ca | 0.4906 |
| ⁵¹V | 0.4945 | ¹²¹Sb | 0.7487 |
| ⁵²Cr | 0.3723 | ¹³⁷Ba | 0.8001 |
| ⁵⁵Mn | 0.4104 | ¹⁸²W | **0.0171** |
| ⁵⁶Fe | 0.6306 | ²⁰⁵Tl | 0.6865 |
| ⁵⁹Co | 0.2228 | ²⁰⁸Pb | **0.0481** |
| ⁶⁰Ni | 0.3475 | ²³²Th | 0.6353 |
| ⁶³Cu | **0.0120** | ²³⁸U | **0.0066** |
| ⁶⁶Zn | 0.5222 |  |  |

Summary of particle-phase metals (ICP-MS) in WS vs FA. These data are from our prior characterization (Wardhani et al. 2024) using the identical generator/chamber settings and are re-presented here for reference. **Bold P values** denote elements with significant elevation in WS at α = 0.05; other elements showed non-significant upward trends.

**Supplementary Table S3. Number of cells per cluster for each biological replicate**

| **Cluster Name** | **FA** | | | **WS** | | |
| --- | --- | --- | --- | --- | --- | --- |
|  | **Rep 1** | **Rep 2** | **Rep 3** | **Rep 1** | **Rep 2** | **Rep 3** |
| Memory CD8+ T cells | 449 | 550 | 499 | 432 | 377 | 368 |
| Neutrophils | 2072 | 2535 | 2115 | 1663 | 1429 | 1339 |
| ISG expressing immune cells | 519 | 669 | 514 | 224 | 280 | 199 |
| Pro-B cells | 362 | 418 | 450 | 288 | 178 | 251 |
| Erythroid-like cells | 1927 | 1739 | 1584 | 1327 | 829 | 1067 |
| Pre-B cells | 826 | 924 | 905 | 554 | 575 | 491 |
| Plasma B cells | 338 | 333 | 403 | 295 | 234 | 278 |
| Naive B cells | 1433 | 1708 | 1598 | 1015 | 1244 | 954 |
| Granulocytes | 78 | 77 | 66 | 40 | 23 | 23 |
| Myeloid Dendritic cells | 143 | 179 | 210 | 163 | 139 | 125 |
| Plasmacytoid Dendritic cells | 173 | 201 | 231 | 148 | 84 | 126 |
| Platelets | 45 | 41 | 39 | 45 | 22 | 24 |
| Basophils | 128 | 138 | 151 | 101 | 78 | 119 |
| Intermediate monocytes | 434 | 567 | 493 | 365 | 319 | 320 |
| Macrophages | 76 | 84 | 76 | 56 | 44 | 33 |
| CD8+ NKT-like cells | 197 | 246 | 212 | 199 | 196 | 138 |
| Total Cells | 9200 | 10409 | 9546 | 6915 | 6051 | 5855 |

This table provides the raw cell counts for each immune cell cluster (rows) for each individual biological replicate (mouse, columns). These absolute counts are the basis for the relative percentage calculations presented in **Figure 3A** and the statistical analyses presented in **Supplementary Figure S3** and **Supplementary Table S4**. 'Total Cells' indicates the total number of quality-controlled cells profiled for each replicate.

**Supplementary Table S4. Statistical results from mixed-effects logistic regression for scRNA-seq cell compositional changes.**

| Cell Type | Estimate | P-Value | Odds Ratio |
| --- | --- | --- | --- |
| * Memory CD8+ T cells | 0.20832086 | 0.0000002 | 1.23160829 |
| * ISG expressing immune cells | -0.4670778 | 0.0000044 | 0.62683134 |
| Neutrophils | 0.0282786 | 0.4417310 | 1.02868224 |
| Pro-B cells | -0.1151142 | 0.2829717 | 0.89126435 |
| Erythroid-like and erythroid precursor cells | -0.0756803 | 0.5456982 | 0.92711258 |
| Pre-B cells | -0.0611774 | 0.1837954 | 0.94065631 |
| Plasma B cells | 0.15783356 | 0.0646364 | 1.17097128 |
| Naive B cells | 0.05862229 | 0.5532800 | 1.06037466 |
| * Granulocytes | -0.5091914 | 0.0000611 | 0.60098131 |
| * Myeloid Dendritic cells | 0.22386417 | 0.0172793 | 1.25090109 |
| Plasmacytoid Dendritic cells | -0.0980374 | 0.4471260 | 0.90661503 |
| Platelets | 0.11627693 | 0.4464050 | 1.12330691 |
| Basophils | 0.10111189 | 0.3829086 | 1.10640043 |
| Intermediate monocytes | 0.04274717 | 0.3273490 | 1.04367399 |
| Macrophages | -0.1368655 | 0.2081670 | 0.87208755 |
| * CD8+ NKT-like cells | 0.2368055 | 0.0012608 | 1.26719463 |

This table provides the full statistical output for the mixed-effects logistic regression model visualized in **Supplementary Figure S3B**. This analysis was performed to robustly validate the cell compositional changes (summarized in **Figure 3A**) between the Wood Smoke (WS) and Filtered Air (FA) groups, accounting for inter-animal variability (n=3 mice/group).

**Supplementary Table S5. Transcriptional downregulation of M2-polarizing regulators in the bone marrow of OVX mice after WS exposure**

| **Gene**  **Symbol** | **Average Log₂FC**  **(WS vs. FA)** | **Adjusted**  **p-value** | **% Cells Expressing**  **(FA)** | **% Cells Expressing**  **(WS)** |
| --- | --- | --- | --- | --- |
| Cd36 | -0.734 | < 0.0001 | 7.7 | 11.1 |
| Stat6 | -0.248 | < 0.0001 | 10.6 | 13.1 |
| Klf4 | -0.329 | < 0.0001 | 10.1 | 12.5 |
| Irf4 | -0.096 | 0.0008 | 10.7 | 12.5 |

Differential expression analysis results for key M2-polarizing transcription factors and the PPARγ-target gene *Cd36* from the global single-cell RNA-seq data (all immune cells combined). The table shows the average log₂ fold change (calculated as log2(mean expression in WS / mean expression in FA)), Benjamini-Hochberg adjusted p-value, and the percentage of cells expressing each gene in the filtered air (FA) and wood smoke (WS) exposed groups (n=3 mice per group). All listed regulators were significantly downregulated across the bone marrow immune compartment in OVX mice following WS exposure.

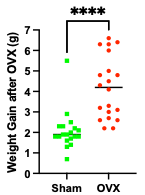

**Supplementary Figure S1. Validation of effective ovarian hormone deficiency by weight gain.**

Change in body mass (Δg) from pre-surgery baseline to the pre-exposure time point for Sham-operated and ovariectomized (OVX) mice used in the functional BMDM assays. OVX mice exhibited significantly greater weight gain compared to Shams, providing physiological validation of effective estrogen deficiency prior to WS/FA exposure. Data are presented as scatter plot with median indicated. Significance was determined by Mann-Whitney U test (**p < 0.0001). N = 19 Sham, N = 20 OVX.

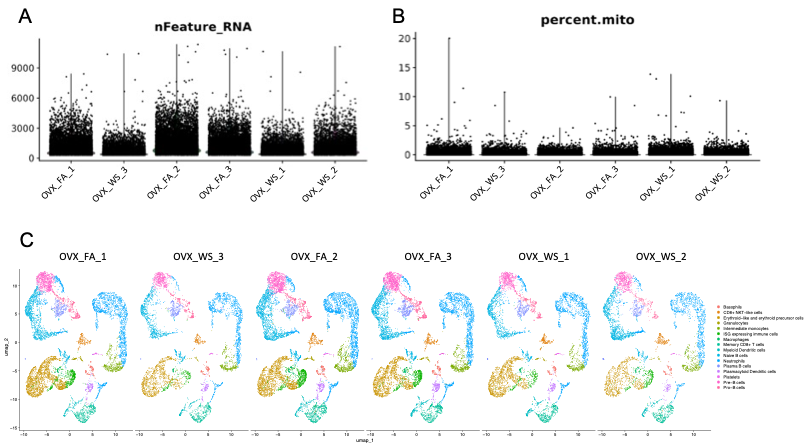

**Supplementary Figure S2. Quality control metrics confirm consistent scRNA-seq sample integrity across groups.**
(A) Violin plot showing the number of detected genes per cell (*nFeature_RNA*) for each biological replicate. (B) Violin plot showing the percentage of mitochondrial gene expression (*percent.mito*) across all cells per replicate. Each distribution represents a single sample from one of the two experimental groups: OVX-FA or OVX-WS (*n = 3 mice per group*). (C) Per-sample UMAP plots for each biological replicate (3-FA, 3-WS), demonstrating consistent cluster identification and data integrity across all samples. All samples passed quality control thresholds and demonstrated comparable sequencing complexity and cellular integrity. See also Figure 1.

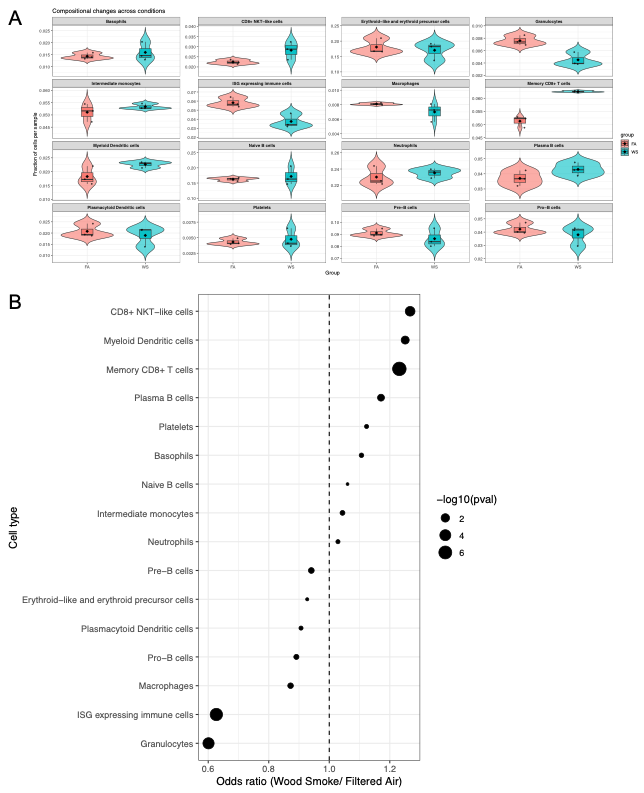

**Supplementary Figure S3. Robust statistical validation of cell compositional changes.**

To confirm the statistical robustness of cell compositional changes observed with n=3 replicates, a mixed-effects logistic regression model was performed. (A) Violin plots showing the fraction of each cell type per individual biological replicate (mouse) for Filtered Air (FA, red) and Wood Smoke (WS, blue) groups. (B) Forest plot summarizing the results of the mixed-effects logistic regression. Dots represent the Odds Ratio (Wood Smoke/Filtered Air) for each cell type. An odds ratio > 1.0 (right of the dashed line) indicates the cell type is more likely to be found in the WS group (increased proportion), while an odds ratio < 1.0 (left of the line) indicates a decreased proportion. The size of the dot corresponds to the statistical significance -log10(*pval*). This robust analysis confirms the significant changes identified in Figure 3A (Memory CD8+ T cells, ISG expressing immune cells, Granulocytes) and reveals additional significant increases in Myeloid Dendritic cells and CD8+ NKT-like cells. Full statistical details, including p-values and 95% confidence intervals, are provided in Supplementary Table S4.

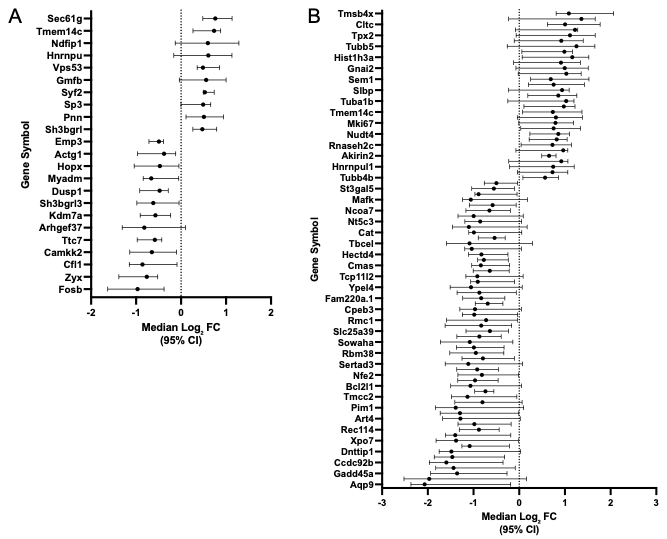

**Supplementary Figure S4. Effect sizes for all remaining significant DEGs from Figure 4 and 5.**

Forest plots display the median log₂FC and 95% confidence intervals (CI) for all significant differentially expressed genes (DEGs) that met the significance criteria (adjusted p < 0.05 and |log₂FC| > 0.5) but were not included in the main figures' core pathway analyses. This figure is provided to ensure full data transparency and completeness. (A) Forest plots for the 23 remaining DEGs (13 down-regulated, 10 up-regulated) from memory CD8⁺ T cells (related to Figure 4). (B) Forest plots for the 87 remaining DEGs (57 down-regulated, 30 up-regulated) from ISG-expressing immune cells (related to Figure 5).

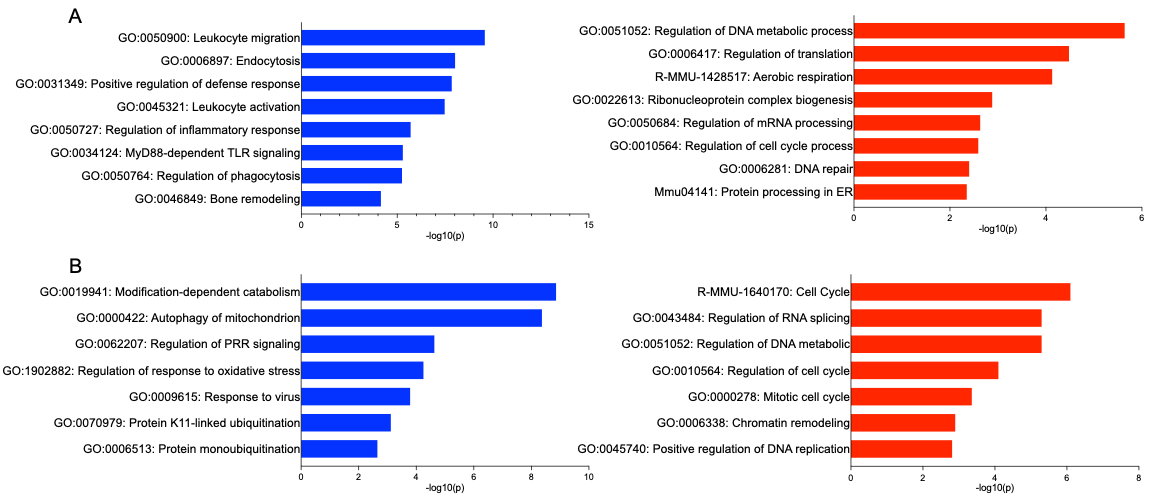

**Supplementary Figure S5. Pseudobulk pathway analysis validates main transcriptional themes at the animal-replicate level.**

To assess animal-level robustness and overcome the low statistical power of n=3 replicates, we performed pathway analysis (Metascape) on pseudobulk DEGs defined by a relaxed statistical threshold (unadjusted p < 0.05, |log2FC| > 0.5). Bar plots show the top curated biological pathways identified from this analysis, selected for relevance to the main findings in Figure 4 and 5. Pathways enriched from downregulated genes are shown in blue, while pathways from upregulated genes are shown in red. The X-axis represents the -Log10(P) value, indicating statistical significance. The full Top 20 pathway lists for this analysis are provided in Supplementary Table S6. (A) Memory CD8+ T cells. The analysis confirmed the significant suppression of key immune function pathways (e.g., leukocyte migration, leukocyte activation) and the activation of stress-response pathways (e.g., regulation of DNA metabolic process, DNA repair). This result clearly validates the core themes identified in the cell-level analysis (Figure 4B, 4D). (B) ISG-expressing cells. The analysis similarly confirmed the suppression of protein regulation and anti-viral pathways (e.g., modification-dependent protein catabolism, response to virus) and the strong activation of genomic stress pathways (e.g., Cell Cycle, chromatin remodeling). This result clearly validates the core themes identified in the cell-level analysis (Figure 5B, 5D).

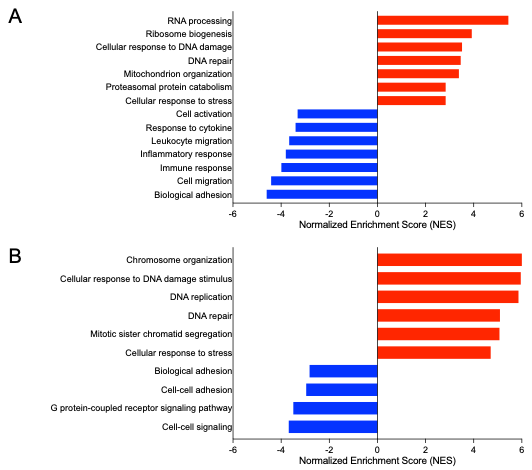
**Supplementary Figure S6. GSEA provides orthogonal validation for core transcriptional themes.**

Bar plots show the Normalized Enrichment Score (NES) for key curated GO Biological Process pathways identified by GSEA. Pathways enriched from downregulated genes are shown in blue, while pathways from upregulated genes are shown in red. This analysis, which uses the entire ranked gene list, provides independent, orthogonal validation for the biological themes identified in our main DEG pathway analysis (Figure 4 and 5). (A) In Memory CD8⁺ T cells, GSEA confirms the suppression of pathways related to immune activation and migration (validating Figure 4B) and the strong activation of pathways related to cellular stress, DNA repair, and RNA processing (validating Figure 4D). (B) In ISG-expressing cells, GSEA confirms the overwhelming activation of pathways related to genomic stress, cell cycle, and DNA repair (validating Figure 5D) and the suppression of pathways related to cell-cell signaling and adhesion (complementing Figure 5B). All pathways shown are highly significant (FDR < 0.01). Full GSEA results are provided in Supplementary Table S7.

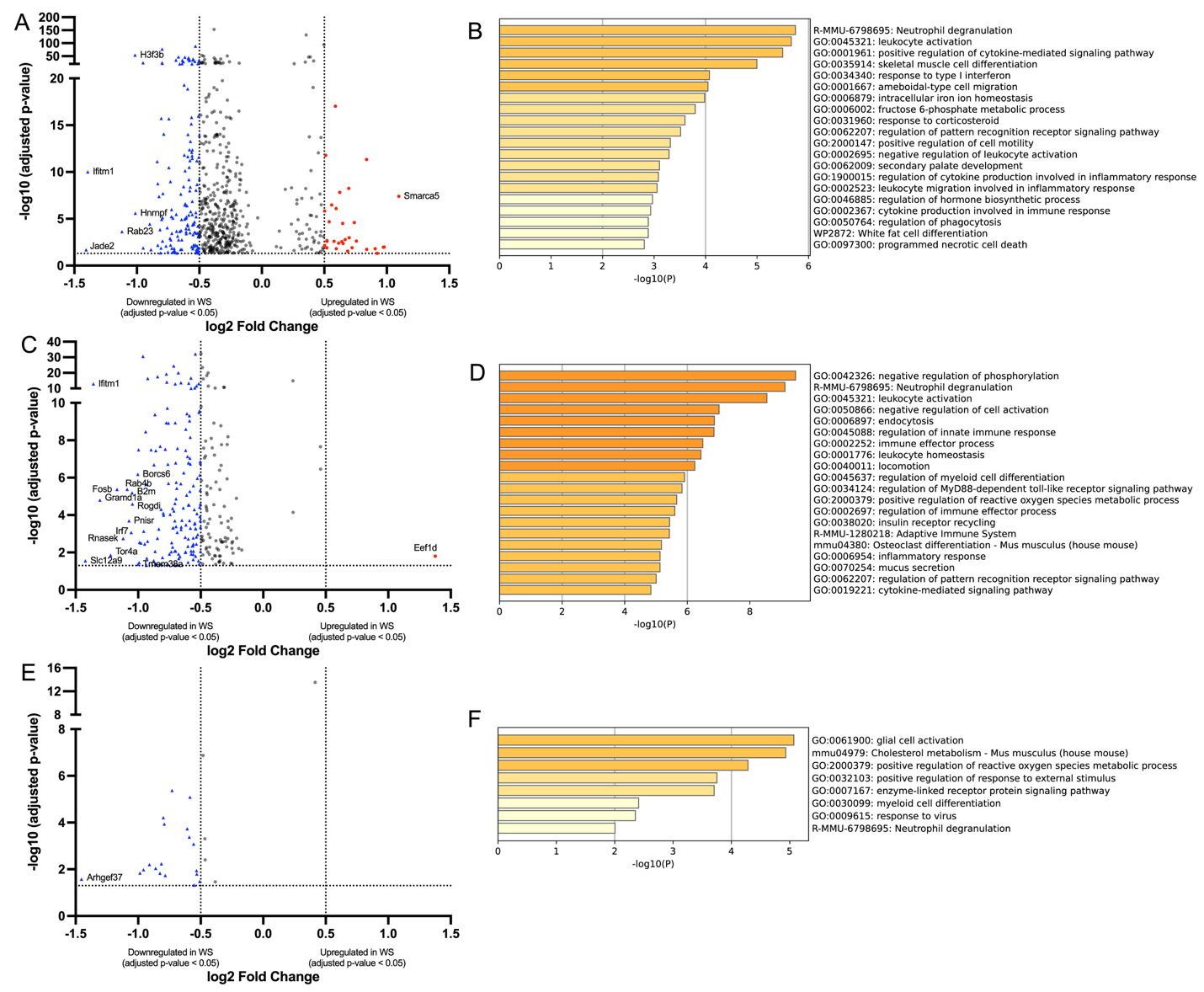

**Supplementary Figure S7. Wood smoke suppresses innate immune pathways in neutrophils, monocytes, and dendritic cells in OVX bone marrow**

Volcano plots (A, C, E) show differentially expressed genes (DEGs) in neutrophils (A), intermediate monocytes (C), and dendritic cells (E) from OVX mice exposed to WS versus FA. Significantly upregulated (red circles) and downregulated (blue triangles) genes are highlighted, respectively (adjusted p < 0.05 with |log₂FC| > 0.5). Bar graphs (B, D, F) display top enriched biological pathways for downregulated genes in neutrophils (B), monocytes (D), and dendritic cells (F). The color intensity of each bar corresponds to its p-value significance, with darker shades indicating more significant pathways. WS exposure led to transcriptional suppression in pathways associated with neutrophil degranulation, cytokine signaling, leukocyte migration, and type I interferon response. n = 3 biological replicates per group were used for scRNA-seq, and DEGs were identified by the Wilcoxon rank-sum test with false discovery rate correction. See also Figure 3, Supplementary Figure S3, and Table S4.

and Figure S1.

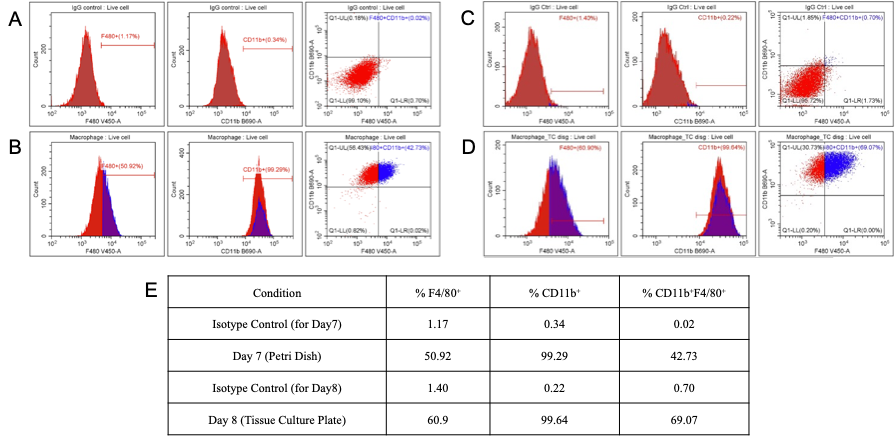

**Supplementary Figure S8. Flow cytometric validation of BMDM maturation before and after re-plating on tissue culture plate.**

Representative flow cytometry analysis of bone marrow cells following 7 days of M-CSF differentiation and subsequent re-plating. (A) Analysis of cells harvested directly from the differentiation petri dish (Day 7), showing high CD11b expression (>99%) but intermediate F4/80 expression (~51% F4/80⁺, ~43% CD11b⁺F4/80⁺). Percentages shown are relative to the corresponding Day 7 isotype controls. (B) Analysis of cells after re-plating onto tissue culture plates and adhering for 24 hours (Day 8, conditions used for functional assays), showing sustained high CD11b expression (>99%) and a marked increase in the mature, double-positive (CD11b⁺F4/80⁺) population (~69%). Percentages shown are relative to the corresponding Day 8 isotype controls. (C) Tabular summary quantifying the percentages of F4/80⁺, CD11b⁺, and CD11b⁺F4/80⁺ cells among live singlets for both conditions and their respective isotype controls (data from one representative experiment, n=3). The consistently high CD11b⁺ percentage confirms a highly enriched myeloid/macrophage lineage. The increase in the CD11b⁺F4/80⁺ fraction from Day 7 (differentiation culture) to Day 8 (on assay plate) reflects adherence-associated maturation typical of *in vitro* M-CSF derived BMDMs.

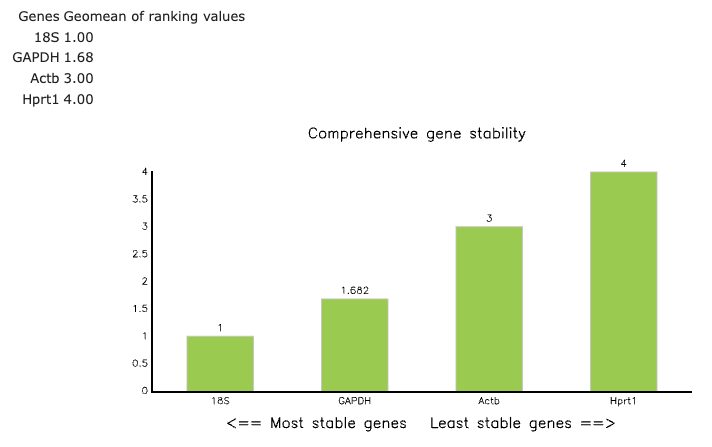

**Supplementary Figure S9. Validation of reference gene stability across experimental conditions.**

To empirically determine the most stable reference gene for qPCR normalization, the expression of four candidate genes (18S, GAPDH, Actb, and Hprt1) was measured across all experimental samples (n=5 per group). The stability of these genes was evaluated using the RefFinder tool, which integrates multiple algorithms to generate a comprehensive stability ranking. The graph displays the final stability value (Geomean of ranking values) for each gene, where a lower value indicates higher expression stability. The analysis unequivocally identified 18S rRNA as the most stable reference gene in our experimental system.

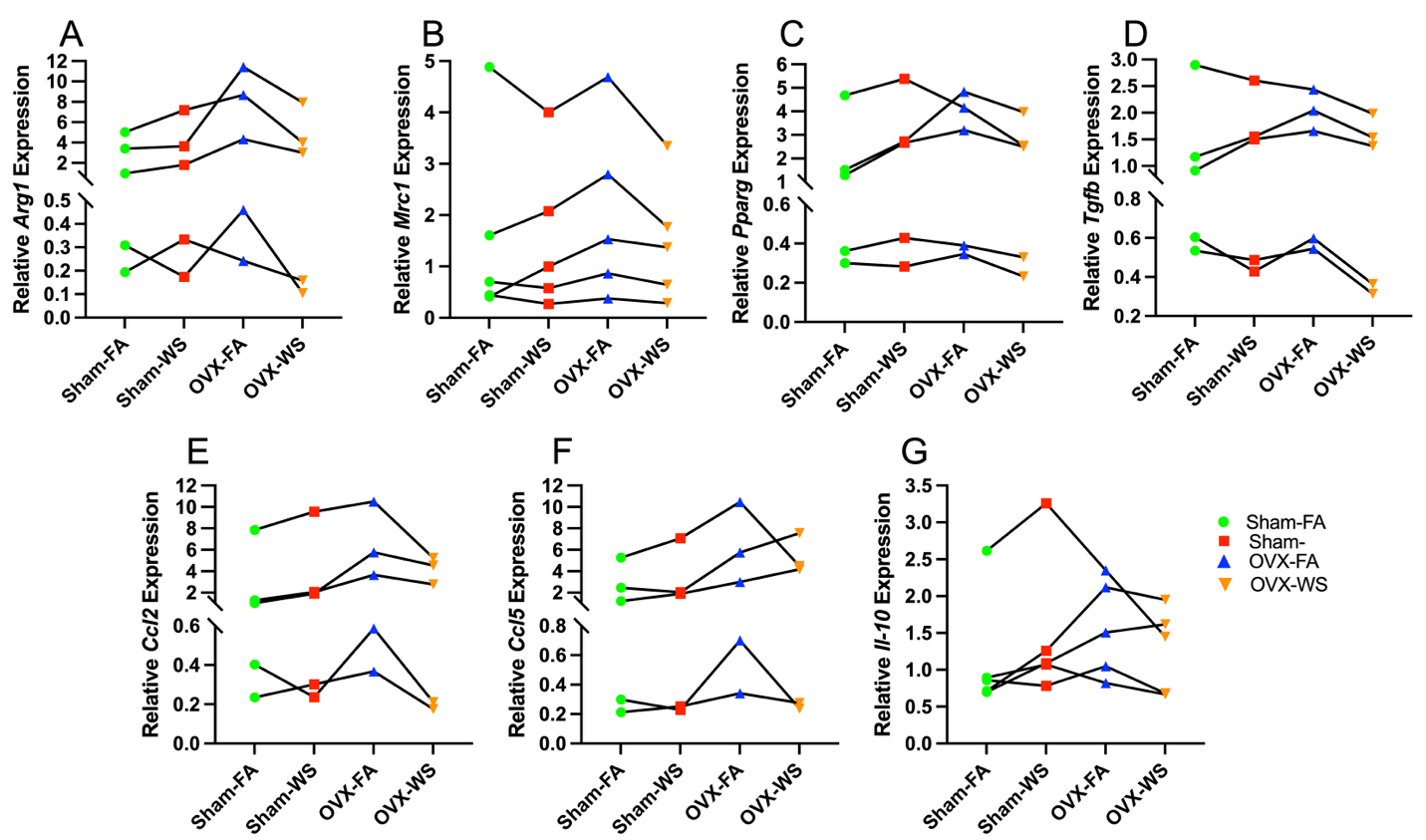

**Supplementary Figure S10. Individual response patterns of M2-associated genes show consistent downregulation by wood smoke in OVX-derived macrophages.**

(A–G) Profile plots for the relative mRNA expression of seven M2-associated genes: (A) *Arg1*, (B) *Mrc1*, (C) *Pparg*, (D) *TGgfb*, (E) *Ccl2*, (F) *Ccl5*, and (G) *Il10.*  Data shown are the same relative quantification values (2−ΔΔCt) as presented in Figure 7. Each line represents a single biological replicate (n=5), and the data points were connected to visualize the response pattern across the experimental groups. These plots provide the visual justification for employing a linear mixed-effects model for the analysis in Figure 7. They clearly illustrate both the significant inter-replicate variability (i.e., batch effects, evidenced by the different baseline levels of each line) and the consistent within-replicate response patterns that are appropriately accounted for by modeling 'Biological Replicate' as a random effect (See Figure 7).

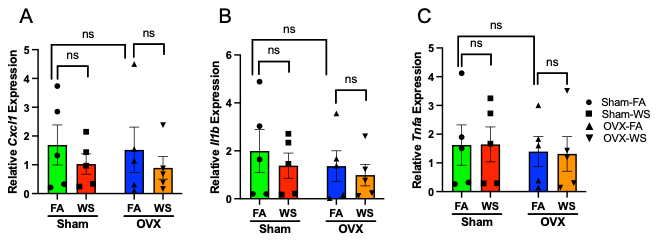

**Supplementary Figure S11. Expression of M1-associated genes is not significantly altered by WS in Sham or OVX BMDMs**

(A–C) Bar graphs show relative mRNA expression of M1-associated genes *CXCL1* (A), *IL-1β* (B), and *TNF-α* (C) in bone marrow–derived macrophages (BMDMs) from Sham and OVX mice exposed to FA or WS. For visualization, gene expression levels were normalized to 18S rRNA and relative quantification was calculated using the 2−ΔΔCt method, with the mean ΔCt of the Sham-FA group serving as the calibrator for all samples. Bars represent mean ± SEM for each group, and each data point represents an individual mouse (n = 5 biological replicates per group). Statistical significance was determined using a linear mixed-effects model (LMM) on the ΔCt values. No significant main effects or interaction effects were observed across groups. ns = not significant (See Figure 7).
